# *Notch* expression during ctenophore development gives insight into its ancestral function

**DOI:** 10.1101/2025.09.30.679519

**Authors:** Brent Foster, Fredrik Hugosson, Cezar Borba, James Strother, Mark Q. Martindale

**Affiliations:** The Whitney Laboratory for Marine Bioscience, Department of Biology, University of Florida, St. Augustine, FL 32080, USA

**Keywords:** Notch, molecular evolution, ctenophores

## Abstract

The canonical *Notch* pathway is a juxtacrine signaling module with widely conserved roles maintaining progenitor cell populations, promoting binary cell fate decisions, and establishing tissue boundaries. This pathway emerged in Metazoan lineages, although many molecular components and regulators existed prior to that divergence. Given that the vast majority of *Notch* studies focus on bilaterians, it is unclear when or how *Notch* gained its developmental signaling functions. To clarify the ancestral function of *Notch*, we turned to an early branching Metazoan — the ctenophore *Mnemiopsis leidyi* — and conducted structural analyses of putative *Notch* components to evaluate predicted signaling potential. We characterized gene expression of these components with *in situ* hybridization and show they are expressed during late embryogenesis. Using hybridization chain reaction (HCR^TM^), we examine *MlNotch* expression relative to canonical transcriptional targets and putative stem cell markers and identify differential expression patterns at a cellular resolution. Pharmacological inhibition reveals zones of active and inactive *MlNotch* during late embryonic development. Our results suggest that *Notch* evolved its signaling potential by the Metazoan divergence and is likely involved in regulating progenitor cell populations and coordinating developmental fate decisions.

## Introduction

The canonical *Notch* pathway is a juxtacrine signaling module involved in various aspects of development across Metazoa. It is best known for its role in promoting binary cell fate decisions via lateral inhibition and/or induction, maintaining stem and progenitor cell populations, and regulating terminal differentiation (for reviews of the role of *Notch* during development, see ^1,2^). Many of the key regulatory enzymes and transcriptional targets for *Notch* signaling predate the divergence of Metazoan lineages, suggesting that this pathway evolved by co-opting molecules from pre-existing signaling modules^3,4^ (see also Figure 1A).

**Figure 1.**
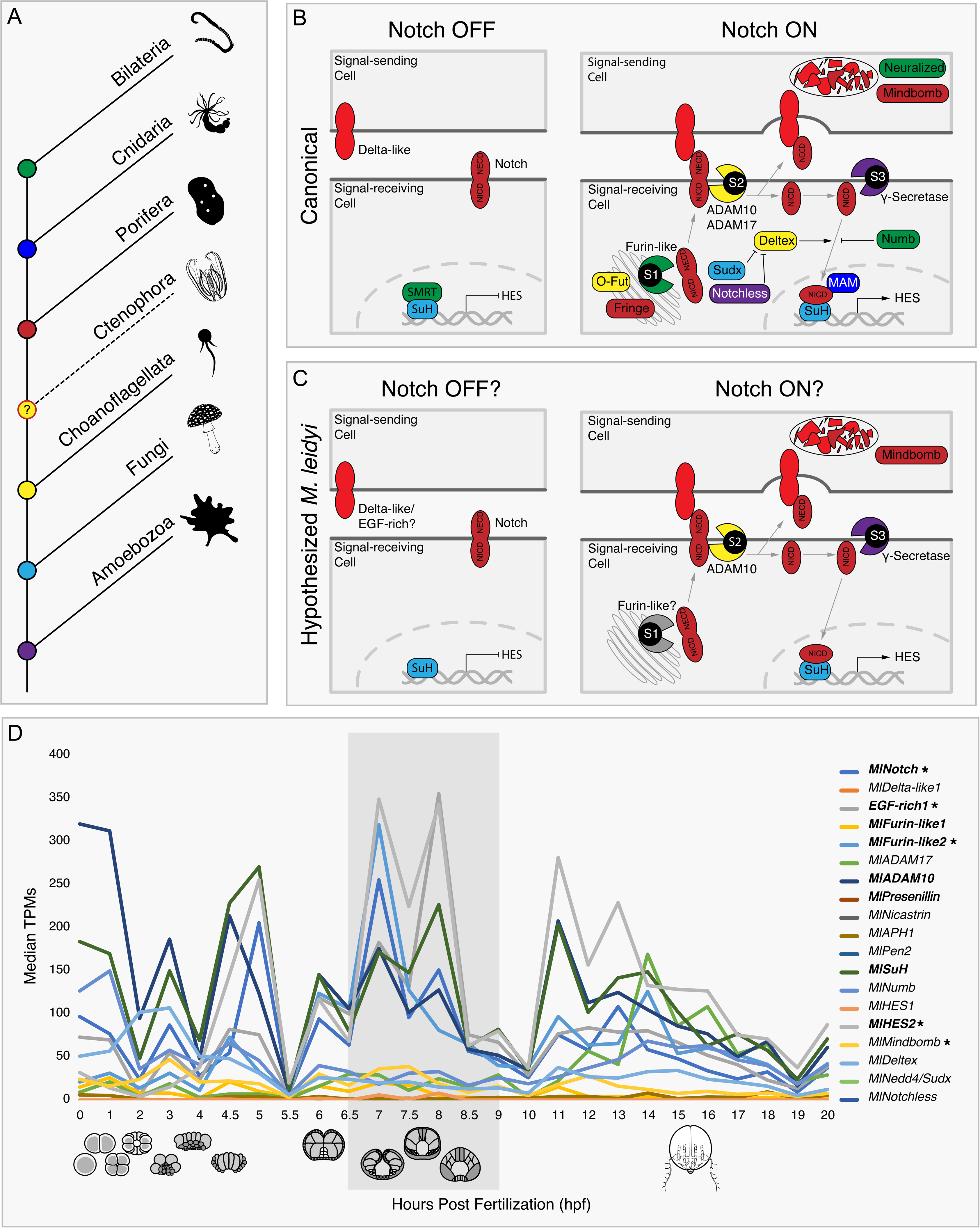
The canonical *Notch* signaling pathway emerged in Metazoan lineages. A) Simple phylogeny depicting when molecular components of the canonical *Notch* pathway evolved. B) Cell diagram of canonical *Notch* pathway in bilaterians, with components color-matched to A) when they are believed to have first emerged. C) Hypothetical cell diagram of components of the putative canonical Notch pathway encoded in the *Mnemiopsis leidyi* genome. D) Developmental RNA-seq of putative canonical *Notch* pathway genes^37^. Bolded genes in the key represent genes expressed during the developmental peak shaded in gray. * = previously unreported genes.

Canonical *Notch* signaling consists of a transmembrane protein that undergoes several distinct proteolytic cleavages for its activation. In the Golgi apparatus, Furin-like convertase forms a *Notch* heterodimer (S1 cleavage) that translocates to the cell membrane, where it acts as a signal receptor. The *Notch* receptor can then interact with a ligand expressed in a neighboring cell, exposing a second extracellular cleavage site recognized by a membrane-bound metalloprotease (S2 cleavage). This cleavage releases the *Notch* extracellular domain (NECD) from the membrane, allowing it to be taken up by the neighboring signaling cell. Meanwhile, γ-secretase in the signal-receiving cell recognizes a third cleavage site and releases the *Notch* intracellular domain (NICD) from the membrane (S3 cleavage), whereupon it translocates to the nucleus and acts as a co-factor to regulate transcription (see Figure 1B for a detailed cell diagram depicting canonical *Notch* signaling in bilaterians).

The canonical *Notch* pathway is likely a Metazoan innovation and is present in virtually all bilaterian model systems, although when and how this signaling module emerged during animal evolution remains a mystery. Genomic evidence in single-celled Eukaryotes suggests that domain shuffling of separate proteins could have led to the formation of Metazoan *Notch*^5,6^. A more recent report has identified a *Notch-like* gene with conserved PFAM domains in their canonical order in the choanoflagellate species *Mylnosiga fluctuans*, although to date no study has identified a non-Metazoan species that encodes both *Notch* receptor and a recognizable activating ligand^7^. The vast majority of *Notch* studies are conducted in bilaterians and have demonstrated a broad conservation of components and functions^4,8–10^. Comparative genomics has shown that *Notch* orthologs and putative pathway components have been identified in each non-bilaterian clades, however, to our knowledge *in situ* characterizations of *Notch* components in non-bilaterians have only been conducted in a handful of sponges^11,12^ and cnidarians^13–17^. To better understand how the *Notch* pathway and its various developmental functions evolved, it is necessary to understand the developmental context in which these components are expressed in other early branching Metazoan representatives.

Ctenophores, as the likely sister group to all other Metazoans^18–20^, can offer key evolutionary insight into how *Notch* signaling and its developmental functions emerged in Metazoans, regardless of whether a canonical pathway is intact or not. *Mnemiopsis leidyi,* the primary subject of this report, encodes at least 16 out of 22 canonical *Notch* pathway components in its genome, with several other components reported to be missing so-called diagnostic domains, raising questions concerning their orthology and role in a functional pathway^18^. Existing data only provide presence or absence information which has led subsequent reports to equate the absence of these canonical *Notch* components and PFAM domains as evidence for the absence of a functional signaling pathway^9,10,18^. To date, no study has analyzed the cellular and developmental context in which putative canonical components are expressed during ctenophore development. We therefore have examined the predicted signaling capacity of *M. leidyi Notch* (*MlNotch*) and the spatiotemporal expression of putative pathway regulators and targets during early ctenophore development to gain insight into how these components may have evolved to form a functional developmental signaling module.

## Results

### M. leidyi genome encodes many canonical Notch components with characteristic PFAM domains

Many components of the canonical *Notch* pathway are definitively encoded in the genome of the ctenophore *Mnemiopsis leidyi*^18^, although these characterizations have been limited to an absent/present dichotomy that assumed that missing components or diagnostic domains of putative *Notch* genes equated non-functionality. However, if these putative canonical components are expressed in the same or adjacent cells at the same time, they could ostensibly behave as a rudimentary pathway (compare Figure 1B to Figure 1C). We therefore examined the cellular and developmental context in which these components are expressed to test the hypothesis that canonical *Notch* components are expressed in shared domains during early ctenophore development and function as a primitive pathway.

**Figure S1.**
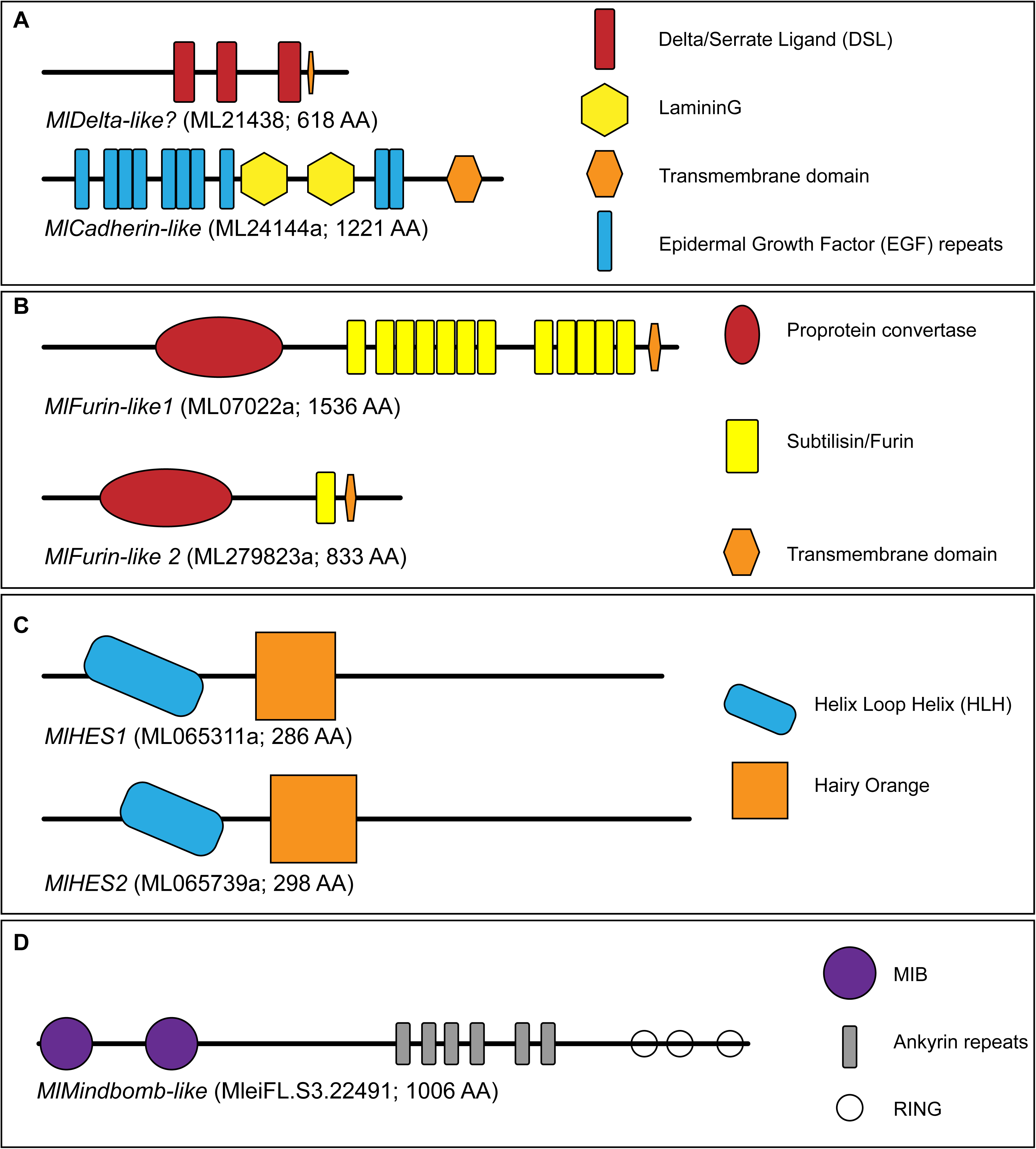
Domain architectures of proteins implicated in the canonical *Notch* pathway. A) Candidate ligands, *MlDelta-like* and *MlCad-like*. B) Putative S2 proteases, *MlFurin-like1* and *MlFurin-like2*. C) Hairy-Enhancer of Split orthologs, *MlHES1* and *MlHES2*. D) E3 ubiquitin ligase, *MlMindbomb-like*.

We first confirmed that these ctenophore *Notch* components contain all of their characteristic PFAM domains. We conducted independent BLAST searches of canonical *Notch* components against the *M. leidyi* genome and confirmed hits with reciprocal BLAST and PFAM analysis. Upon examination of the reported *MlDelta-like* (ML21438a) gene, we were able to identify the delta-serrate ligand (DSL) PFAM domains previously reported missing (see Figure S1A and File S1). However, reciprocal BLAST searches did not result in any hit with matching PFAM domains, leading us to conclude that this hit was an error in the genome sequencing. Furthermore, BLAST searches of the *Drosophila Delta-like* sequence against the *M. leidyi* genome yielded the gene identified as *MlNotch* (ML128617a) and a gene rich in both EGF repeats and LamininG domains, characteristic of proto-cadherins (ML24144a; see Figure S1A and File S1). Given that these genes lack both DSL and MNNL domains, we do not consider them true *Delta-like* proteins. We therefore refer to ML24144a as *MlCadherin-like* (*MlCad-like*). Our searches also revealed that the Furin-like gene previously reported^18^ (ML07022a) retains several Furin-like repeats and subtilase domains characteristic of Furin convertases; we therefore refer to this gene as *MlFurin-like1*. We also identified previously unreported orthologs for a second Furin-like convertase (ML279823a; Figure S1B and File S1), a second HES transcription factor (ML065739a; Figure S1C and File S1), and an ortholog of the *Delta-like* regulator *Mindbomb* (MleFL.S3.22491; Figure S1D and File S1). We refer to these genes as *MlFurin-like2*, *MlHES2*, and *MlMindbomb*, respectively.

Using existing developmental RNA-seq data from *M. leidyi*^21^, we plotted the median transcripts per million (TPMs) of the putative *Notch* components encoded in the genome identified in our independent searches. We noted three apparent peaks of gene expression at approximately 4 hours post fertilization (hpf), 8hpf, and 12hpf (Figure 1D). We opted to focus our gene expression studies on the 8hpf timepoint to avoid potential confounds with the maternal-to-zygotic transcriptional transition^22^. The genes expressed in this 8hpf peak correspond to putative components of the core canonical *Notch* pathway: *MlNotch* (receptor), *MlPCad-like* (candidate ligand), *MlFurin-like2* (S1 cleavage), *MlADAM10* (S2 cleavage), *MlPresenilin* (S3 cleavage), *MlSuH* (co-factor), and *MlHES2* (transcriptional target).

### Notch negative regulatory regions and intracellular domains are highly conserved

We next wanted to evaluate the conserved signaling capacity of the *Notch* receptor by conducting a phylogenetic analysis of *Notch* orthologs across Metaozoa and analyzing their respective PFAM domain architectures and juxta-transmembrane domains (J/TM). Lin-*Notch* Repeats (LNR) and Ankyrin repeats characteristic of *Notch* evolved before the Metazoan lineages. In fact, the LNR and *Notch* Intracellular Domain (NICD) are remarkably conserved across Metazoans, whereas the extracellular Epidermal Growth Factor (EGF) repeats are highly variable (Figure 2A). Our phylogram of *Notch* has strong support placing ctenophore *Notch* proteins at the base of Metazoa, supporting previous supposition that NICD architecture was present in the last common animal ancestor^7^. While each of the predicted ctenophore *Notch* proteins contains conserved nuclear localization signals, we were unable to identify these domains in all of the representative holozoan *Notch*-*like* proteins.

**Figure 2.**
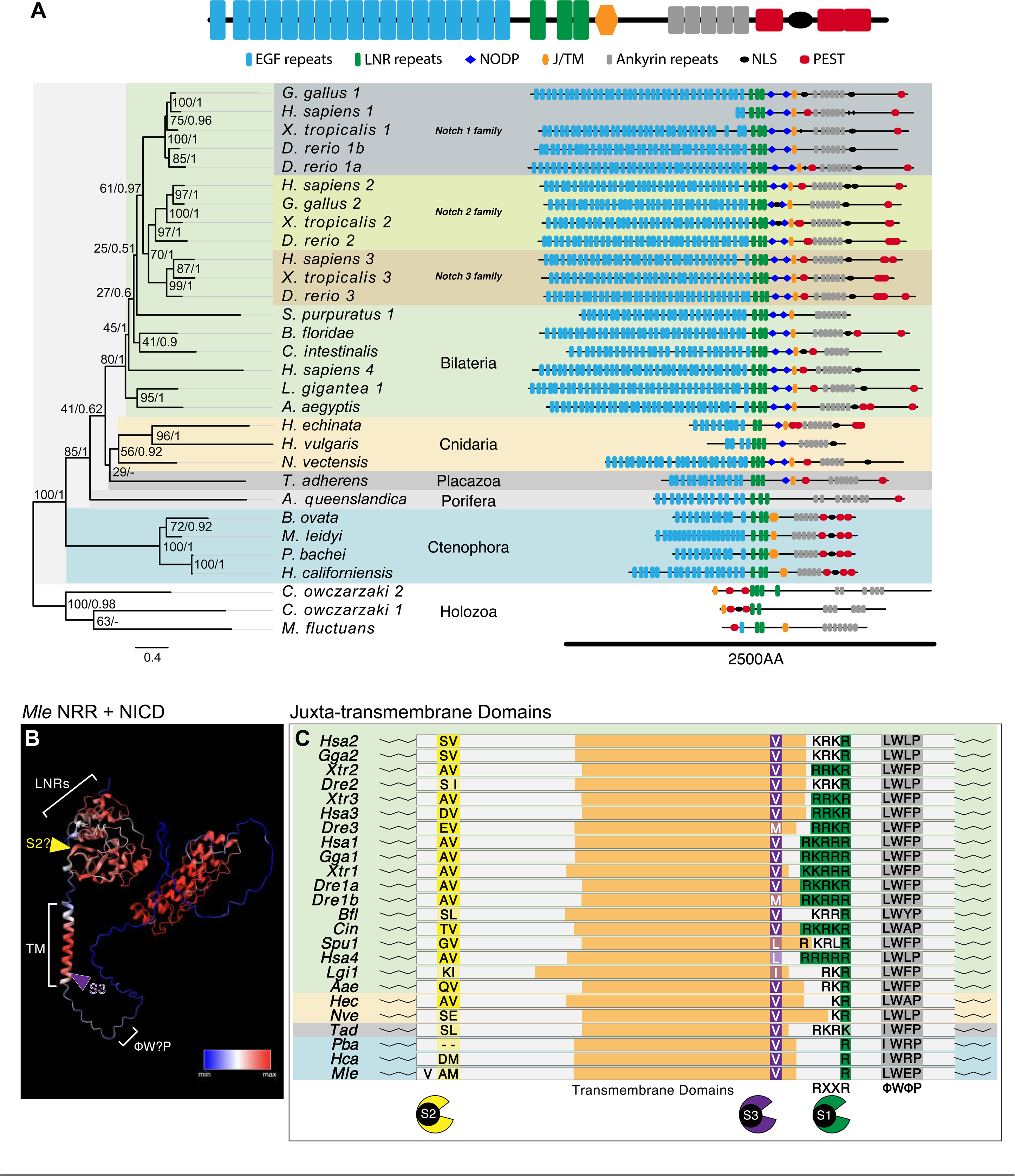
*Notch* domains predate Metazoans. A) Maximum likelihood/Bayesian phylogeny of *Notch* across Metazoa showing conserved *Notch* Intracellular Domains (NICDs). Node labels are Maximum Likelihood / Posterior Probabilities values. B) AlphaFold 3D prediction of *MlNotch* Negative Regulatory Region (NRR) and NICD. C) Basic alignments of *Notch* juxta-transmembrane domains highlighting conserved S1, S2, and S3 cleavage recognition sites. Species in panel C) are color-matched to phyla from panel A).

Given that canonical *Notch* signaling activity requires three discrete cleavage events, we wanted to evaluate whether *MlNotch* has conserved regulatory structures and cleavage recognition sites. Our AlphaFold prediction shows that *MlNotch* LNRs cover the extracellular J/TM region in *MlNotch* (Figure 2B). Our multiple sequence alignment shows that residues recognized by Furin-like and γ-secretase are highly conserved, whereas the putative S2 cleavage site is poorly conserved (Figure 2C). The minimal Furin-like recognition motif (RXXR) is only observed in bilaterian lineages, although we found a highly conserved Arginine residue aligned in nearly all of the examined sequences. W and P residues of the ΦWΦP motif are also highly conserved. Taken together, these analyses suggest that *MlNotch* may retain signal-transducing capacity independent of S1 or S2 cleavage via Furin-like convertase and ADAM metalloproteases, respectively.

### Putative Notch components share expression domains during late ctenophore embryogenesis

We examined gene expression of *MlNotch* and *MlCad-like* during late embryonic development. *MlNotch* transcripts are broadly expressed at 8hpf in developing pharynx, tentacle bulbs, and aboral organ and in ectodermal cells invaginating at the oral pole (Figure 3A). We also identified a peculiar symmetrical *MlNotch* expression pattern in groups of ectodermal cells at the aboral end of developing embryos (Figure 3A’). *MlCad-like* is expressed in the developing pharynx and tentacle bulbs, similar to *MlNotch* (Figure 3B and Figure 3B’), although we failed to detect clear expression in the aboral end (Figure 3B’). We did not detect expression of the previously identified *MlDelta-like* 8hpf (Figure S2). All four putative protease genes share broad expression domains at 8hpf in the developing pharynx and tentacle bulbs, although we failed to detect expression of these genes in the aboral end (Figure 3C–3F).

**Figure 3.**
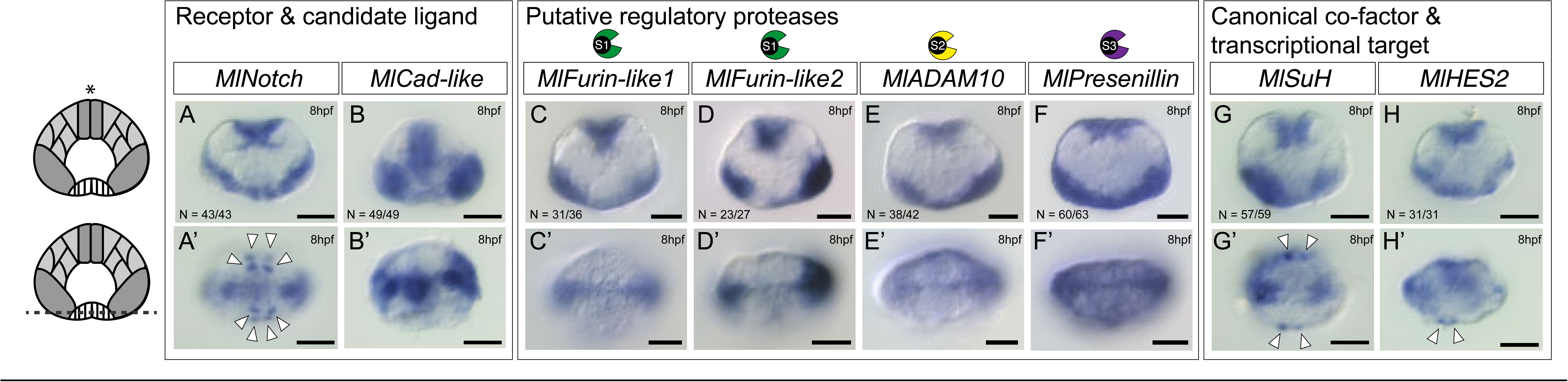
Genes implicated in canonical *Notch* signaling are expressed in developing pharynx and tentacle bulbs at 8hpf. A–A’) *MlNotch*. B–B’) *MlCad-like*. C–C’) *MlFurin-like1*. D–D’) *MlFurin-like2*. E–E’) *MlADAM10*. F–F’) *MlPresenillin*. G–G’) *Suppressor-of-Hairless (MlSuH)*. H–H’) *MlHairy-Enhancer-of-Split2 (MlHES2)*. * = oral pole. All scalebars = 50µm. White arrowheads indicate distinct patches of expression in aboral cells.

**Figure S2.**
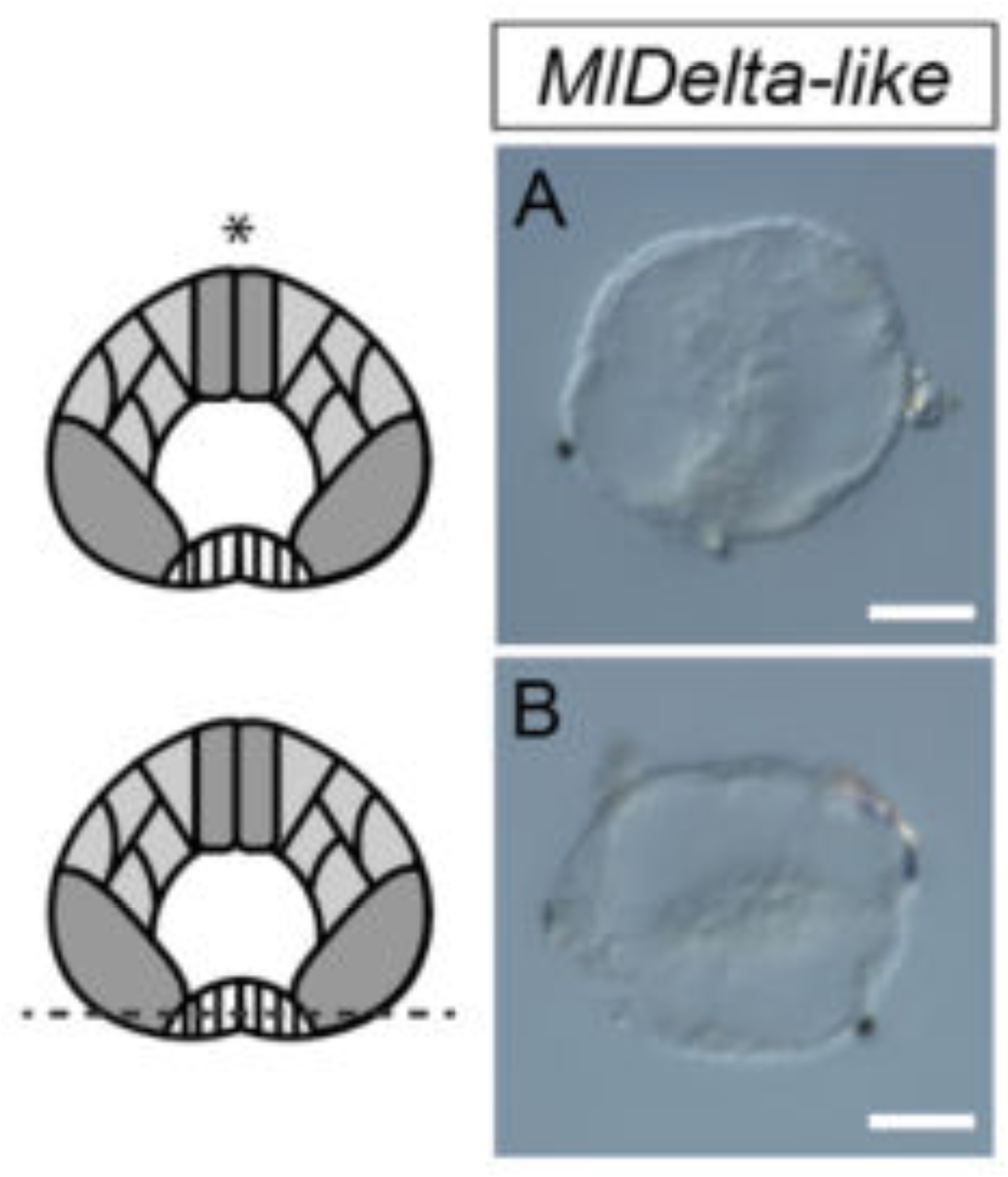
*MlDelta-like* is not detectable 8hpf. A) Lateral view. B) Aboral view. * = oral pole. All scalebars = 50µm.

We next examined gene expression of a putative *MlNotch* co-factor (*MlSuH*) and transcriptional target (*MlHES2*). Colorimetric *in situs* of *MlSuH* illustrate a similar expression pattern to *MlNotch* with broad expression domains in the developing pharynx and tentacle bulbs (Figure 3G). We also note patches of *MlSuH* expression in the aboral end of the developing embryo, similar to our observations for *MlNotch* (Figure 3G’). *MlHES2* is expressed in similar domains (Figure 3H and Figure 3H’).

### MlNotch and a candidate ligand are expressed in distinct cell populations

To obtain a more cellular resolution of these gene expression patterns, we modified a hybridization chain reaction (HCR^TM^) protocol^23^. We observed weak expression of *MlNotch* in the developing pharynx and tentacle bulbs and identified a strong *MlNotch* signal in distinct patches of cells at the aboral end of 8hpf embryos (Figure 4A). *MlCad-like* signal is most pronounced at the oral pole and developing tentacle bulbs (Figure 4B). While we note apparent regions where *MlNotch* and *MlCad-like* are co-expressed (Figure 4C; see insert 4C’), we also identify differential expression in the developing pharynx and tentacle bulbs (Figure 4C; see inserts 4C’’–4C’’’). Aboral views confirm the symmetrical nature of *MlNotch* expression in distinct patches of cells, similar to what we observed in our colorimetric *in situs* (Figure 4D; see insert 4D’). *MlCad-like* does not appear to be expressed in the same or neighboring cells in the aboral end of developing embryos (Figure 4E; see insert 4E’). We note regions where *MlNotch* and *MlCad-like* appear to be co-expressed in the ectoderm of the developing tentacle bulbs (Figure 4D–4F; see inserts 4D’’, 4E’’, and 4F’’).

**Figure 4.**
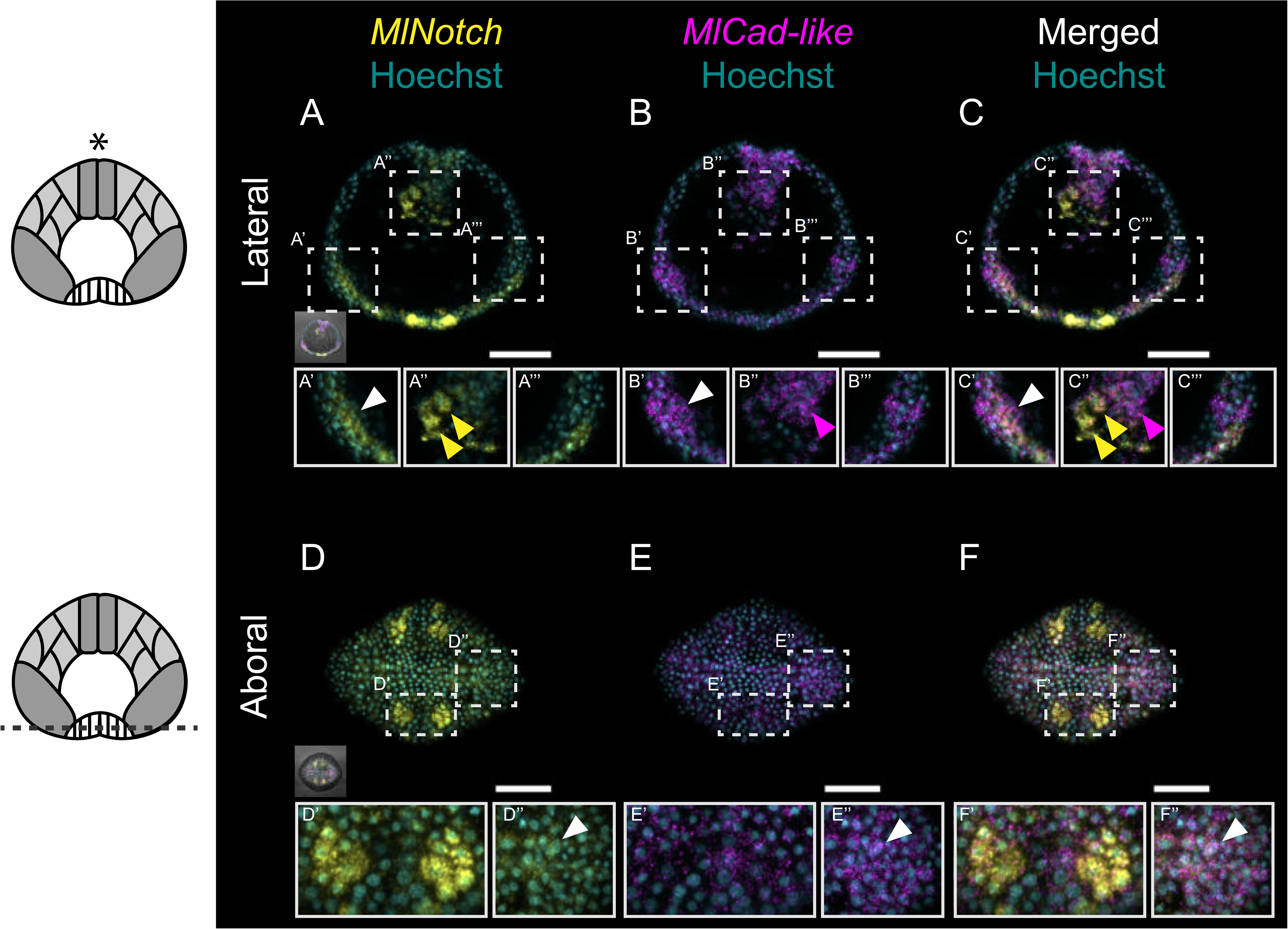
*MlNotch* and *MlCad-like* transcripts are both differentially and co-expressed 8hpf. A–C) Lateral views of *MlNotch* (A), MlCad-like (B), and Merged channels (C). D–F) Aboral views of *MlNotch* (D), *MlCad-like* (E), and Merged channels (F). * = oral pole. All scalebars = 50µm. White arrows indicate cells co-expressing *MlNotch* and *MlCad-like*. Yellow arrows indicate *MlNotch-*positive cells. Magenta arrows indicate *MlCad-like-*positive cells.

### MlNotch-positive cells co-express canonical transcriptional targets or are found adjacent to cells expressing target transcripts

We next considered expression of *MlNotch* relative to *MlHES2* and *MlSox1*, both canonical transcriptional targets. *MlHES2* shares similar expression domains as *MlCad-like* in the developing tentacle bulbs and pharynx 8hpf, with several cells differentially expressed between *MlNotch-*positive domains in the aboral end (Figure 5A–F). Cells in the developing tentacle bulb appear to co-express *MlNotch* and *MlHES2* transcripts (Figure 5D–F), suggestive of two distinct cell populations or perhaps of a transcriptional transition in cell identity during development.

**Figure 5.**
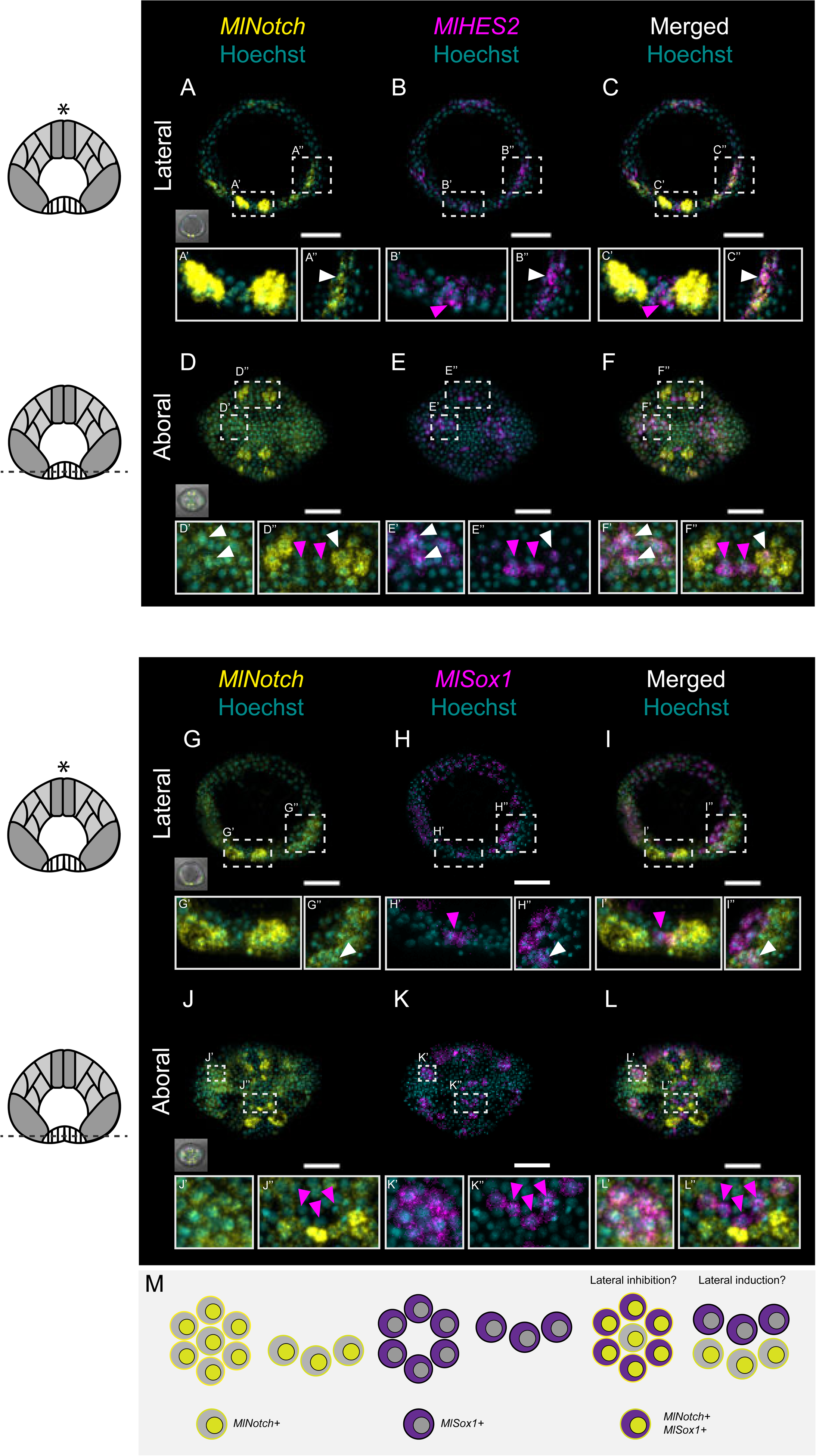
*MlNotch-*positive cells neighbor cells expressing canonical transcriptional targets. A–C) Lateral views of *MlNotch* (A), *MlHES2* (B), and Merged channels (C). D–F) Aboral views of *MlNotch* (D), *MlHES2* (E), and Merged channels (F). G–I) Lateral views of *MlNotch* (G), *MlHES2* (H), and Merged channels (I). Aboral views of *MlNotch* (J), *MlHES2* (K), and Merged channels (L). M) Cartoon schematic representing expression of *MlNotch* relative to *MlSox1* from the panels in J–L. * = oral pole. All scalebars = 50µm. Magenta arrowheads indicate cells expressing *MlHES2* or *MlSox1*, but not *MlNotch*. White arrowheads indicate cells that co-express *MlNotch* and either *MlHES2* or *MlSox1,* respectively.

*MlSox1* is a member of the SoxB gene family and has been shown to be expressed in distinctive domains in the aboral end of developing *M. leidyi* embryos with colorimetric *in situ* hybridization^24^. We corroborate that expression pattern here with HCR^TM^ (Figure 5G–L) and show *MlNotch*-positive cells neighboring *MlSox1-*positive cells in the aboral end and developing tentacle bulbs of embryos 8hpf (Figure 5J–L). We also show a previously unidentified ring of *MlSox1-*positive cells medial to an outer ring of *MlNotch-*positive cells near where the apical organ will develop (Figure 5J–L; see inserts 5J”, 5K”, 5L”). These expression patterns are highly reminiscent of classical models of *Notch*-induced lateral inhibition and lateral induction^1,25,26^ (see Figure 5M).

**Figure S3.**
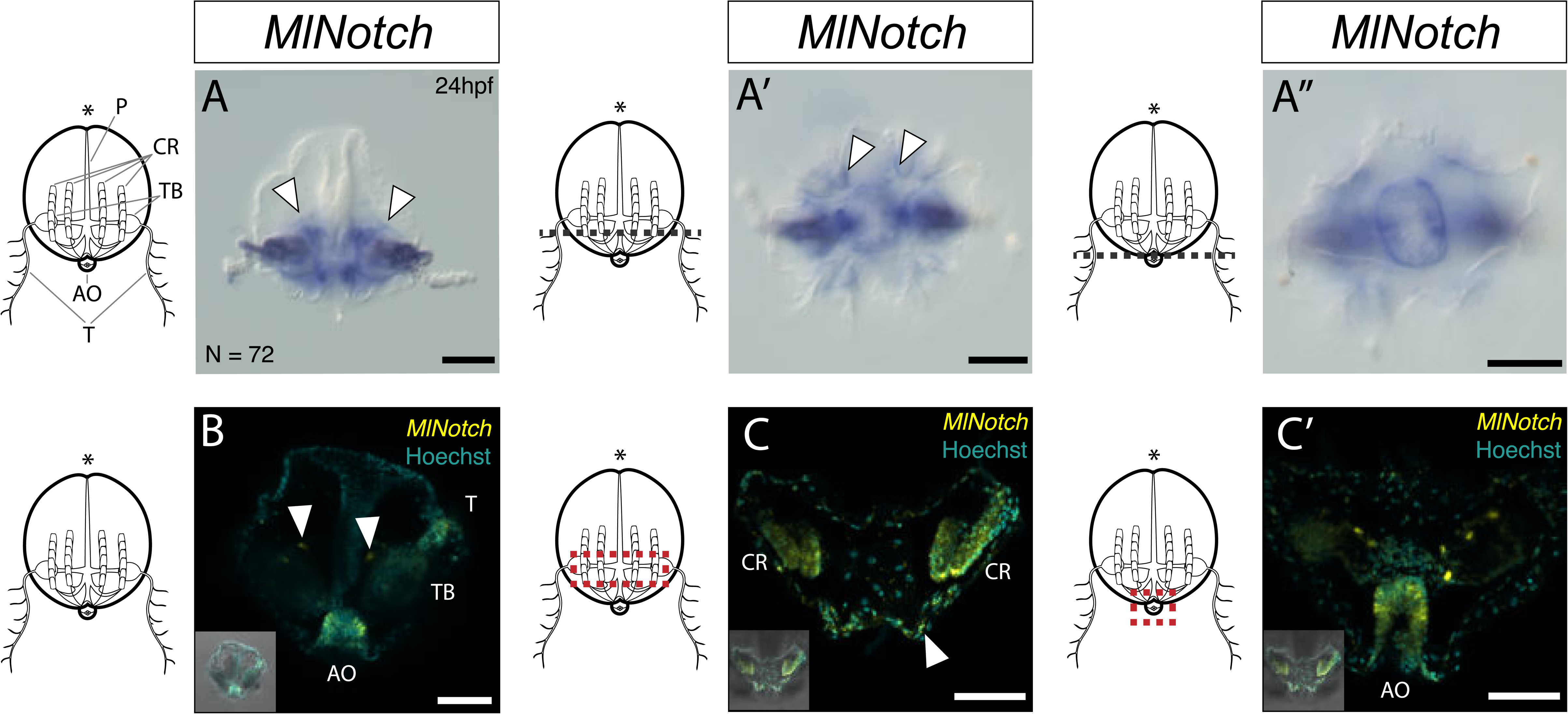
*MlNotch* expression 24hpf. A–A”) Colorimetric ISH of *MlNotch* 24hpf. A) Lateral view showing *MlNotch* expression in mesogleal cells, tentacle bulbs, and floor of the apical organ. A’) Aboral view at the comb row plane showing *MlNotch* expression in presumptive endodermal cells surrounding the comb rows. A”) Aboral view at the plane of the apical organ. B–C’) HCR of *MlNotch* 24hpf. B) Lateral view showing *MlNotch* expression in the floor of the apical organ and in mesogleal cells (see arrowheads). C) Closeup lateral view of *MlNotch*-expressing cells that line the endodermal canals and are sub-adjacent to the comb rows (see arrowhead). C’) Closeup lateral view of *MlNotch-*expressing cells in the floor of the apical organ. * = oral pole. All scalebars = 50µm. P = pharynx, CR = comb rows, TB = tentacle bulb, T = tentacle, AO = apical organ.

### MlNotch is expressed in the floor of the apical organ, comb rows, and endodermal canals 24hpf

We were curious to see how *MlNotch* expression and its implied function might change during ctenophore development. Colorimetric *in situ* hybridization shows *MlNotch* expression in gut regions, tentacles, and the floor of the fully developed apical organ of 24h cydippids (Figure S3A–A”). We also note what appears to be cells in the tissue lining endodermal canals (see arrowheads in Figure S3A and 3A’). Aboral views reveal *MlNotch* expression in cells lining the ciliary comb rows (Figure S3A’) and cells that make up the floor of the apical organ (Figure S3A”). HCR^TM^ experiments corroborate our colorimetric *in situs* with strong *MlNotch* signal in neighboring cells of the apical organ (Figure S3B), comb rows and endodermal canals (Figure S3C and Figure S3C’).

**Figure S4.**
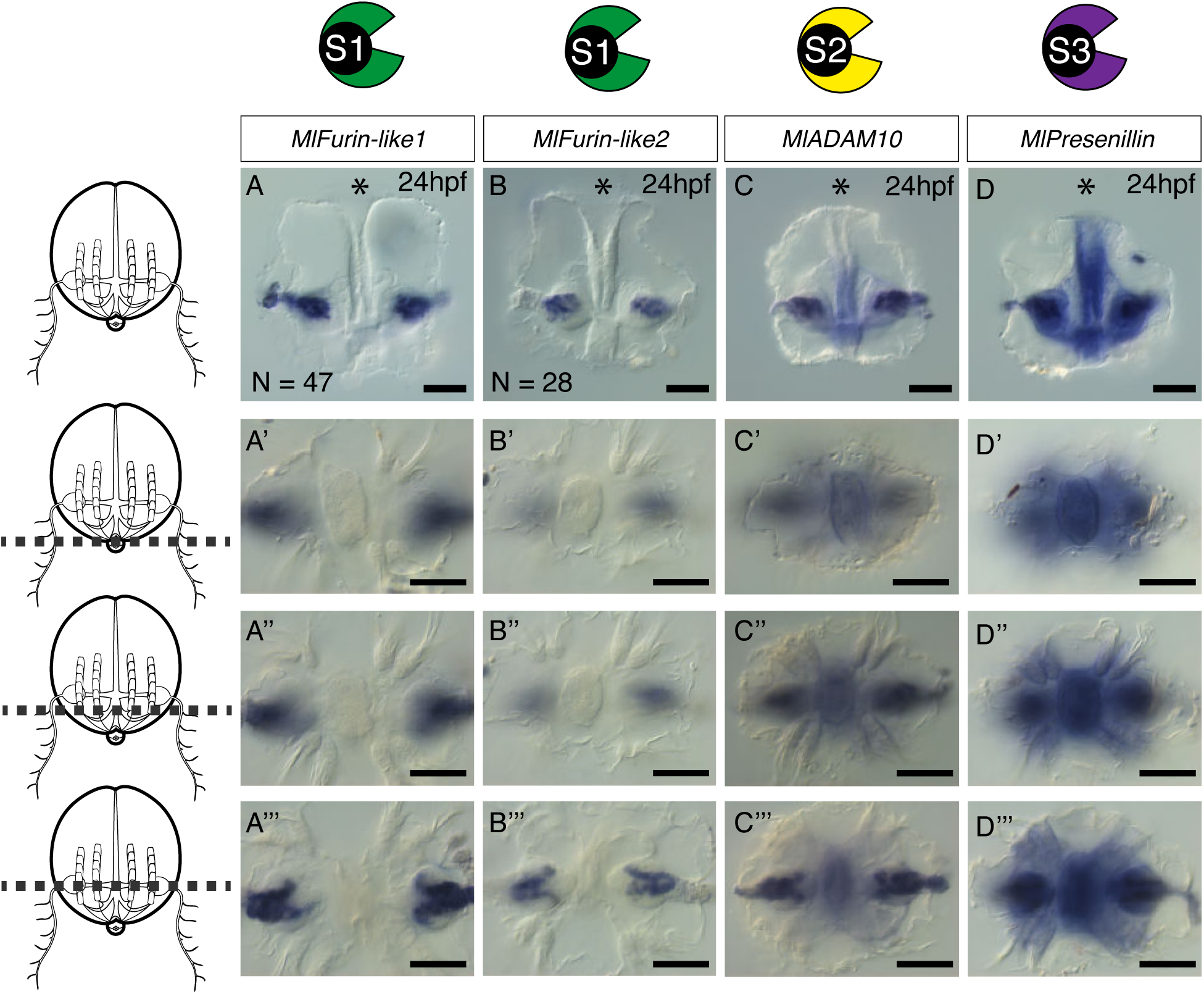
Whole-mount colorimetric *in situ* hybridization of putative genes involved in S1–S3 cleavage. A–A’’’) *MlFurin-like1* expression show in in lateral view (A), plane of the apical organ (A’), plane of the comb rows (A’’), and plane of the tentacle bulbs (A’’’). B–B’’’) *MlFurin-like2* expression show in in lateral view (B), plane of the apical organ (B’), plane of the comb rows (B’’), and plane of the tentacle bulbs (B’’’). C–C’’’) *MlADAM10* expression show in in lateral view (C), plane of the apical organ (C’), plane of the comb rows (C’’), and plane of the tentacle bulbs (C’’’). D–D’’’) *MlPresenillin* expression show in in lateral view (D), plane of the apical organ (D’), plane of the comb rows (D’’), and plane of the tentacle bulbs (D’’’). * = oral pole. All scalebars = 50µm.

### Putative protease genes share expression domains in the apical organ, tentacle bulbs, and pharynx 24hpf

We evaluated expression patterns of putative S1–S3 protease genes at 24hpf via whole-mount colorimetric *in situ* hybridization. Both *MlFurin-like1* and *MlFurin-like2* are exclusively expressed in tentacle bulbs in 24h cydippids (Figure S4A–4B; see also Figure S4A’’’ and 4B’’’). *MlADAM10* is also expressed in the tentacle bulbs and pharynx (Figure S4C) with additional expression in the apical organ (Figure S4C’), around the comb rows (Figure 4SC’’) and tentacle bulb (Figure S4C’’’). *MlPresenillin*, the active subunit of γ-secretase responsible for S3 cleavage, shares similar expression domains in the pharynx, apical organ, and cells surrounding the comb rows (Figure S4D–D’’’). These results support conditions in which these genes may be co-expressed to interact in the tentacle bulbs, although they also hint at differential expression — and therefore function — in cells of the apical organ, comb rows, and pharynx.

### Pharmacological inhibition reveals regions of active and inactive MlNotch

To test whether *MlNotch* is cleaved during late ctenophore embryogenesis, we utilized DAPT and LY411575 — both potent γ-secretase inhibitors^27^ — combined with an antibody that recognizes cleaved *Notch* (see Figure 6A for experimental setup). We note NICD signal localized to the invaginating pharynx and developing tentacle bulbs (Figure 6B), suggesting that these regions are rich in activated *MlNotch*.

**Figure 6.**
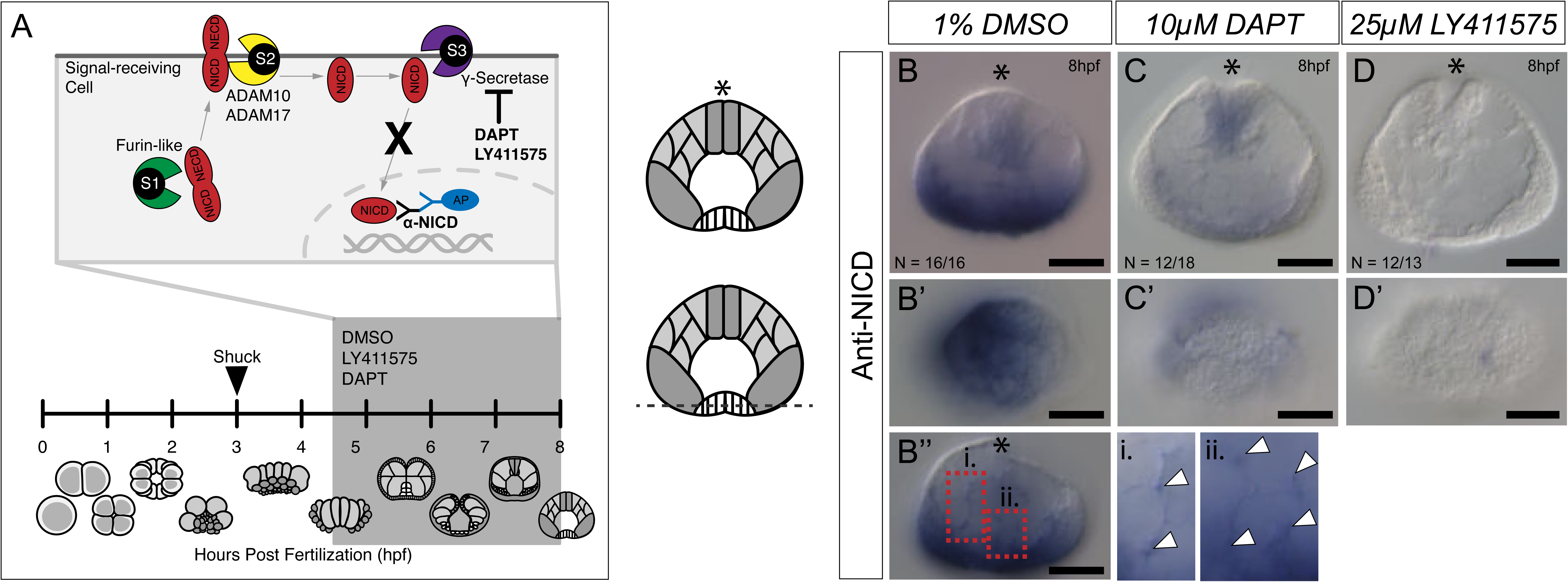
Traditional *Notch* inhibitors reduce expression of *MlNICD* in developing ctenophores. A) Experimental setup of pharmacological inhibition experiments. B–B’’) Representative ctenophore embryo incubated in 1% DMSO. B = lateral view, B’ = aboral view. B’’ shows a lateral view. White arrowheads in i. and ii. point to mesenchymal cells harboring *MlNICD*. C–C’) Representative ctenophore embryo incubated in 10µM DAPT. C = lateral view, C’ = aboral view. D–D’) Representative ctenophore embryo incubated in 25µM LY411575. D = lateral view, D’ = aboral view. * = oral pole. All scalebars = 50µm.

However, we failed to detect signal in the distinctive aboral patches of cells characterized with our colorimetric and HCR *in situs* (Figure 6B’), suggesting that these cell populations are most likely loaded with unprocessed *MlNotch* transcripts or inactive *MlNotch* protein. Mesenchymal cells within the mesoglea also seem to harbor activated *MlNotch* (Figure 6B’’; see white arrow heads). Both DAPT-and LY411575-treated embryos show a noticeable decrease in signal within the pharynx and tentacle bulb regions (Figure 6C & Figure 6D). We did not detect any obvious phenotype or loss of cell types in the embryonic or cydippid stages. Furthermore, drug-treated embryos appear to develop into phenotypically normal cydippids. Together, our results are suggestive of a mechanism where “inactive” *MlNotch* regulates a distinct sub-population of cells that, upon cleavage-induced activation, are expressed in mesenchymal cells and cells that contribute to the future tentacle bulb sheath and pharyngeal epithelium.

## Discussion

### A Notch receptor existed early in the Metazoan divergence and likely retained signaling capacity independent of S1 cleavage

Our bioinformatic structural analysis of the *Notch* juxta-transmembrane domains suggests that *MlNotch* likely does not undergo S1 cleavage. Previous work has identified a Furin-independent function for *Notch* and concluded that this particular role is derived^28,29^, although these studies were limited to cell cultures and *Drosophila*. Here, we show that the minimum Furin-like recognition motif (RXXR) in *Notch* was either lost in ctenophore lineages or a bilaterian innovation. Given that our analysis shows that other non-bilaterian Metazoan *Notch* proteins also lack this exact motif, we favor the latter interpretation. This suggests that S1 cleavage and the heterodimerization of the *Notch* receptor could in fact be derived, which in turn would indicate that the Furin-independent function of *Notch* was ancestral.

Similarly, our analysis shows that the S2 cleavage site seems to be highly diverse in non-bilaterian lineages, although *MlNotch* has an LNR domain architecture that may retain a negative regulatory function by covering an as-of-yet unidentified site recognized by another membrane-bound protease (Figure 2B). Choanoflagellates offer a twist in that their *Notch*-*like* genes also contain structures reminiscent of the negative regulatory regions that, in canonical *Notch* signaling, functions as a cap that covers the S2 sites recognized by ADAM proteinases. In the choanoflagellate *Monosiga brevicolis*, *Notch-like* genes contain an intracellular LNR domain, suggesting that pre-Metazoan *Notch*-*like* proteins may have been capable of intracellular S2-like cleavages, whereas *M. fluctuans* retains the canonical order of PFAM domains and therefore may have been capable of a more canonical extracellular cleavage. Given that *M. brevicolis* is a more recent lineage, this would suggest that the shuffled *Notch* domains is a derived choanoflagellate characteristic and that the pre-Metazoan *Notch* looked — and may have even functioned — similar to the canonical Metazoan *Notch*.

### Developmental and cellular context of MlNotch expression is consistent with models of cell-cell signaling

The putative *Notch* pathway transcripts characterized here all share similar expression domains at the same developmental stage, consistent with what we would expect from components interacting with each other in a functional signaling pathway. The intriguing aboral patches of *MlNotch*-positive cells combined with the data from our pharmacological *Notch* inhibition suggest that these populations are loaded with *MlNotch* transcripts that have not been processed. We hypothesize that these cells are a self-regulating progenitor population characterized by uncleaved — or “inactive” — *MlNotch*. *MlNotch-*positive mesenchymal cells and cells within the developing tentacle bulb and pharynx appear to harbor cleaved — or “active” — *MlNotch*. It is unclear if these active *MlNotch*-positive cells are migrating, but we cannot rule out this possibility. *Notch* signaling has been implicated in coordinating the epithelial-to-mesenchymal transition in cell culture^30^ and regulating migratory peripheral glia in *Drosophila*^31^, motile cells in zebrafish trunk neural crest^32^, and migrating neurons in developing mouse cerebral cortex^33^. To our knowledge, no study has identified a role for *Notch* signaling in regulating migratory cells outside of bilateria. Our results are suggestive of a mechanism where “inactive” *MlNotch* regulates a distinct sub-population of cells that, upon cleavage-induced activation, may be involved in the epithelial-to-mesenchymal transition and the formation of the tentacle bulb sheath and pharyngeal epithelium, pointing to an ancient role for *Notch* regulation of migratory cells. It is unclear what the functions of the other canonical *Notch* pathway genes are in these *Notch-*expressing cells that apparently lack stable cell-cell interactions.

As noted previously, the expression of *MlNotch* in relation to putative transcriptional targets, *MlHES2* and *MlSox1*, is highly reminiscent of classical models of lateral inhibition and induction. This result is intriguing, given that *MlSox1* belongs to the SoxB group^34^ and *Notch* has been shown to have a conserved role in regulating *Sox* genes. Pharmacological inhibition of *Notch* in *Nematostella vectensis* upregulates *SoxB2* genes^35^ while downregulating HES genes^14^. A *Sox*-related gene regulatory network is also moderated by *Notch* in *Hydra vulgaris*^36,37^. *Notch* has also been shown to regulate *Sox* in cephalopod and mouse retinal development^38,39^. *Drosophila* neuroblast formation appears to be gated by a *Notch* and *SoxB* interplay^40^. *SoxB1* coordinates with *NICD* to maintain neural cells in an undifferentiated state in chick embryos^41^. We do not show any clear evidence of *MlNotch* or *MlSox1* promoting neuronal or sensory cell fate in ctenophores, although we do not preclude that possibility. *M. leidyi* embryos develop recognizable neurons by 24hpf, with condensations around the floor of the apical organ by 24hpf^42^, precisely where we see *MlNotch* expressed (Figure S3A’’). Given that *MlNotch* is expressed well before the first presumptive neuron is differentiated, we hypothesize that this embryonic expression is not promoting terminal neuronal fate, although it could very well be involved in regulating early progenitor cell populations. The localization of *MlNotch* expression at 24hpf in endodermal cells subjacent to comb row cells — where the future gonads will develop — hints at a possible role in promoting primordial germ cell fate.

### Pharmacological inhibition of MlNotch points to an as-of-yet unknown mechanism of activation in ctenophores

It is still unclear what activates *MlNotch*. The inability to identify an *MlNotch-*activating ligand has been one of the primary reasons previous reports have concluded that ctenophores do not have a functional *Notch* pathway^9,10,18^. However, the genome of the choanoflagellate species *S. dolichothecata* encodes a protein with MNNL and DSL domains characteristic of *Delta-like*, contrary to previous work suggesting that the *Delta-like* ligand was a Metazoan innovation^7^. Taken together with our ctenophore data, we see at least two possibilities for the emergence of a bonafide *Delta-like* ligand: *Delta-like* could have emerged prior to the Metaozan divergence and was subsequently lost in the ctenophore lineage, or we simply are unable to identify *MlDelta-like* with current methods available to ctenophore researchers.

If *Delta-like* was lost in the ctenophore lineage, then *MlNotch* could either function independent of a ligand or it could recognize an as-of-yet unidentified activating ligand. *Notch* activity can occur during *Drosophila* oogenesis independent of ligand activation^43,44^. Perhaps *MlNotch* is capable of such a ligand-independent function. It may even be possible that the ancestral *Notch* could have functioned as a homodimer, acting as both signal and receiver, and *Delta-like* emerged after the ctenophore divergence. It has been hypothesized that the DSL domains characteristic of *Delta-like* proteins shares ancestry with LNR domains of the *Notch* receptor^3^. Furthermore, the DSL domain is extremely similar to EGF domains — perhaps genes rich in EGF architecture could be viable candidates for identifying non-canonical binding partners. The gene hit from our BLAST searches — ML24144a — is an intriguing candidate given its EGF-rich and LamininG-rich extracellular domains, generally characteristic of cadherins.

Furthermore, upon re-examination of the genomic architecture surrounding ML24144a, we identified a neighboring gene (ML24143a) that appears to harbor the intracellular domain of a proto-cadherin. It is possible that these two genes are misannotated and are in fact a single proto-cadherin that is co-expressed in cells presumably loaded with active *MlNotch*. This raises the intriguing possibility that the ancestral Metazoan *Notch* interacted with cadherin-like proteins. N-cadherins have been shown to interact with Wnt and *Notch* pathways to regulate neural progenitor cells^45^. Our results are consistent with previous hypotheses that *Notch* signaling could have emerged from a cell adhesion module that predates Metazoa^46^. Our HCR^TM^ data show that *MlNotch* and *MlCad-like* transcripts are co-expressed in distinct cell populations. *Delta-like* and *Jagged-like*, both classical *Notch* ligands, have been shown to be *cis-* activators in synthetic cell cultures^47^. It may be possible that *Notch* communication originated as an autocrine signaling module in single-celled eukaryotes that, due to its association with primitive cadherins, evolved its juxtacrine functions at the base of animal multicellularity.

Alternatively, ctenophores could still have a *Delta-like* gene that has evolved such that traditional sequence-based methods are not sensitive enough to identify homologs. Recent work has shown that genes identified as “lineage-specific” may in fact be false-negatives due to the rapid rate in which some genes have evolved such that homologs are unable to be detected^48^. Reasoning that bases homology on sequence similarity or the presence/absence of diagnostic PFAM domains may be similarly flawed. We therefore caution researchers from concluding that specific genes are missing or lineage-specific based on bioinformatic inferences alone, particularly in species where existing genomic and transcriptomic data are prone to errors or mis-annotations.

## Conclusion

This study paints a more complex picture for *Notch* evolution and its activity during ctenophore development than was previously supposed, laying the groundwork to enrich and expand future studies on the evolution of signaling pathways. We have identified and characterized unreported *Notch* pathway components encoded in the *M. leidyi* genome and clarified the developmental context in which these components are expressed. Our results demonstrate that *MlNotch* expression during ctenophore development is consistent with classical models of cell-cell communication, suggesting that its role as a juxtacrine signal transducer may have evolved at the base of the Metazoan divergence, perhaps through a co-option of a pre-existing cell adhesion module. Furthermore, our results highlight the importance of 1) validating bioinformatic data with *in situ* characterizations before making generalized conclusions and 2) expanding the repertoire of model systems to understand the evolution of developmental signaling pathways. While a relative handful of model systems are powerful to study the basic characteristics of signaling pathways, they offer an extremely limited scope for understanding how these pathways evolved. Despite the unique phylogenetic position and intriguing biology of ctenophores, they remain relatively understudied. *In situ* characterizations of signaling pathways in early-branching Metazoans are crucial for understanding how they evolved their developmental functions.

## Acknowledgments

This work was supported by the National Science Foundation (NSF grant IOS-1755364), the National Aeronautics and Space Administration (NASA grant 80NSSC18K1067), and The Paul G. Allen Frontiers Group (grant #12970). The authors thank Dr. Alicia Boyd for her valuable comments while drafting this manuscript.

## Author contributions

Conceptualization, B.F., F.H., and M.Q.M; methodology, B.F., F.H., C.B., and M.Q.M.; validation, B.F., F.H., and C.B.; formal analysis, B.F., F.H., and C.B.; investigation, B.F., F.H., and C.B.; resources, B.F., F.H., C.B., J.S., and M.Q.M; data curation, B.F., F.H., and C.B.; writing – original draft, B.F; writing – review and editing, B.F., F.H., C.B., J.S., and M.Q.M.; visualization, B.F., F.H., and C.B.; supervision, F.H. and M.Q.M.; project administration, J.S. and M.Q.M.; funding acquisition, J.S and M.Q.M.

## STAR Methods

### Resource availability

#### Lead contact

Further information and requests for resources and reagents will be fulfilled by the lead contact, Brent Foster, or Mark Q. Martindale.

#### Materials availability

Animals, reagents, and probes generated by this study are available from Mark Q. Martindale.

#### Data and code availability

This study did not generate any new code.

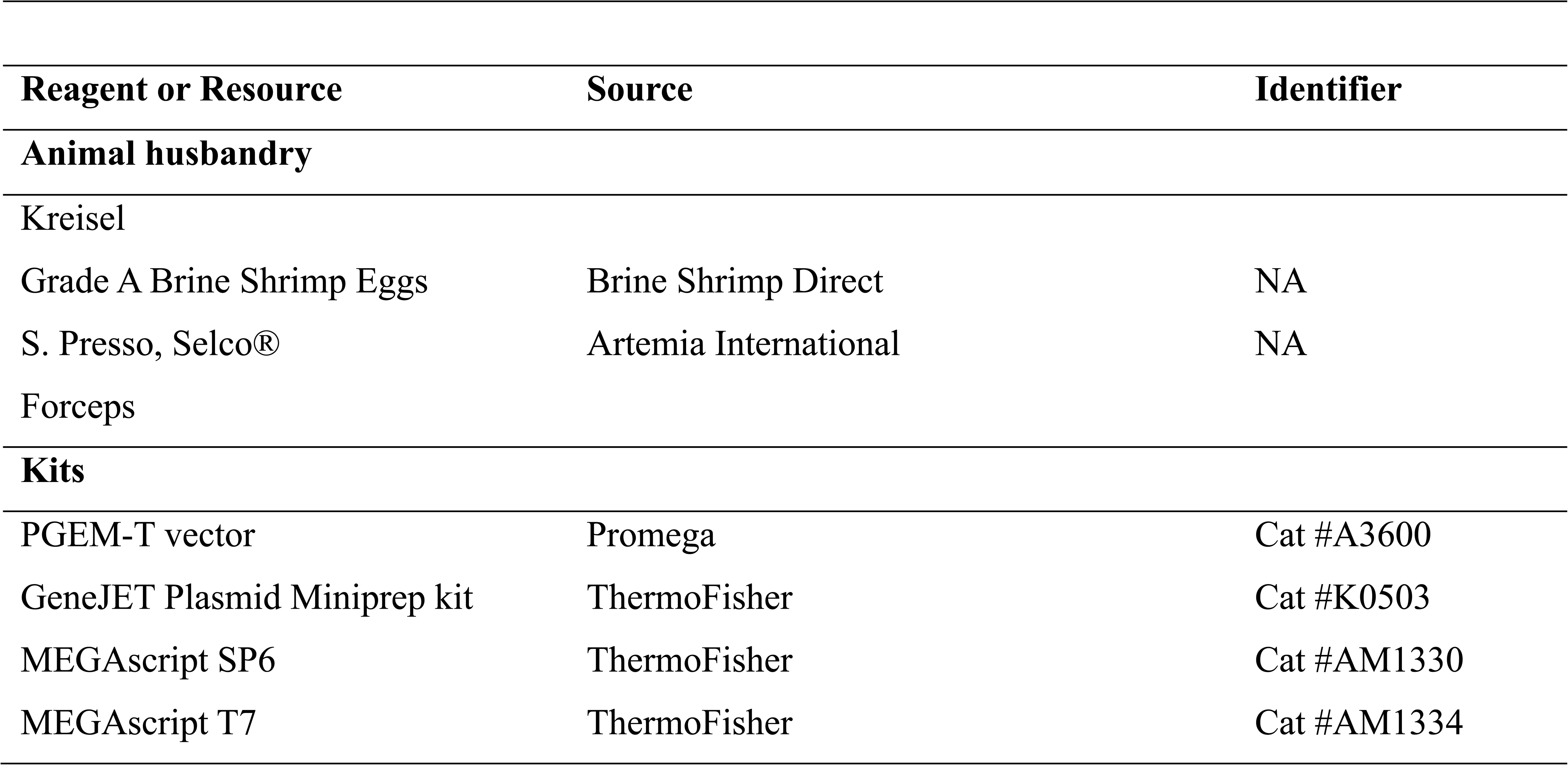
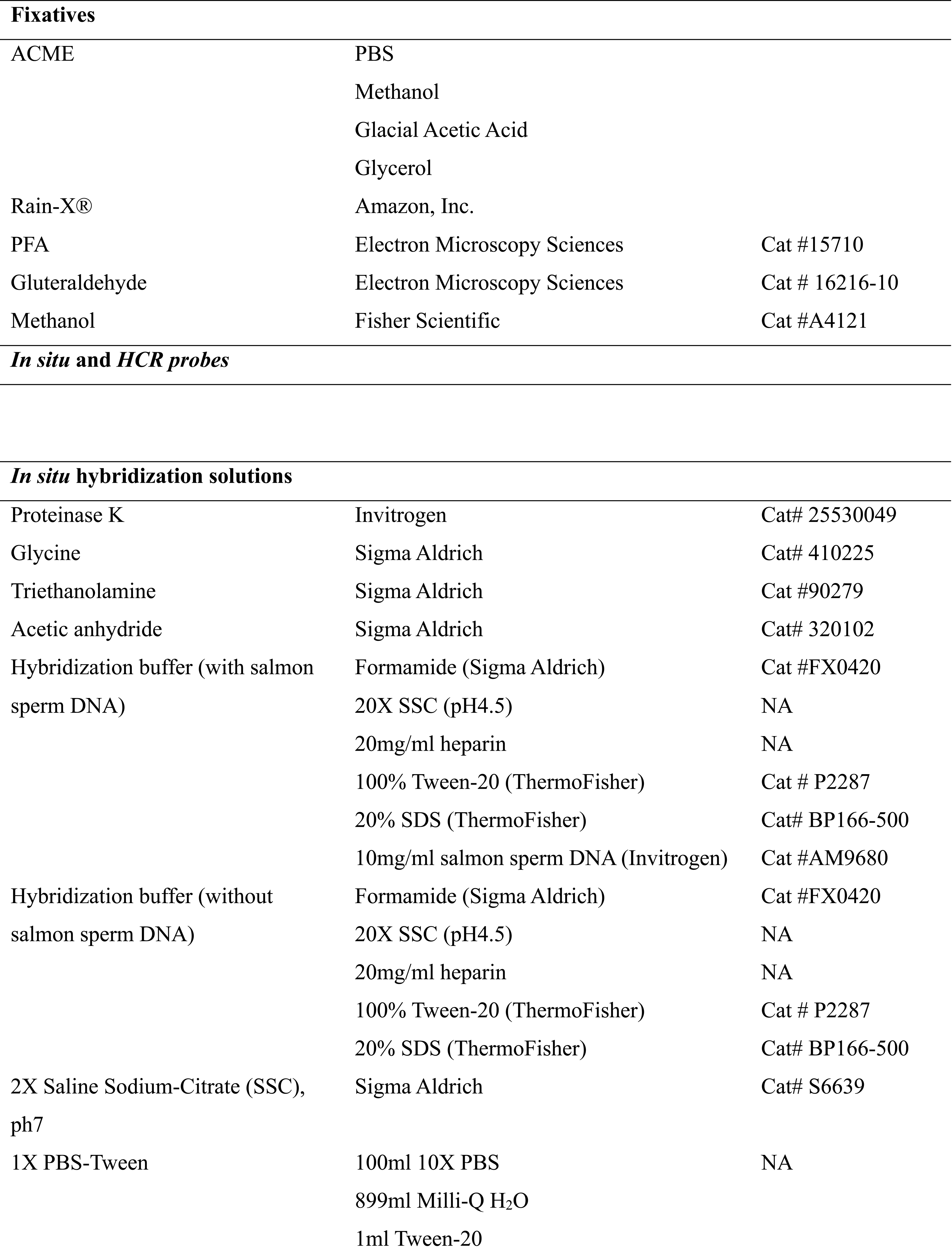
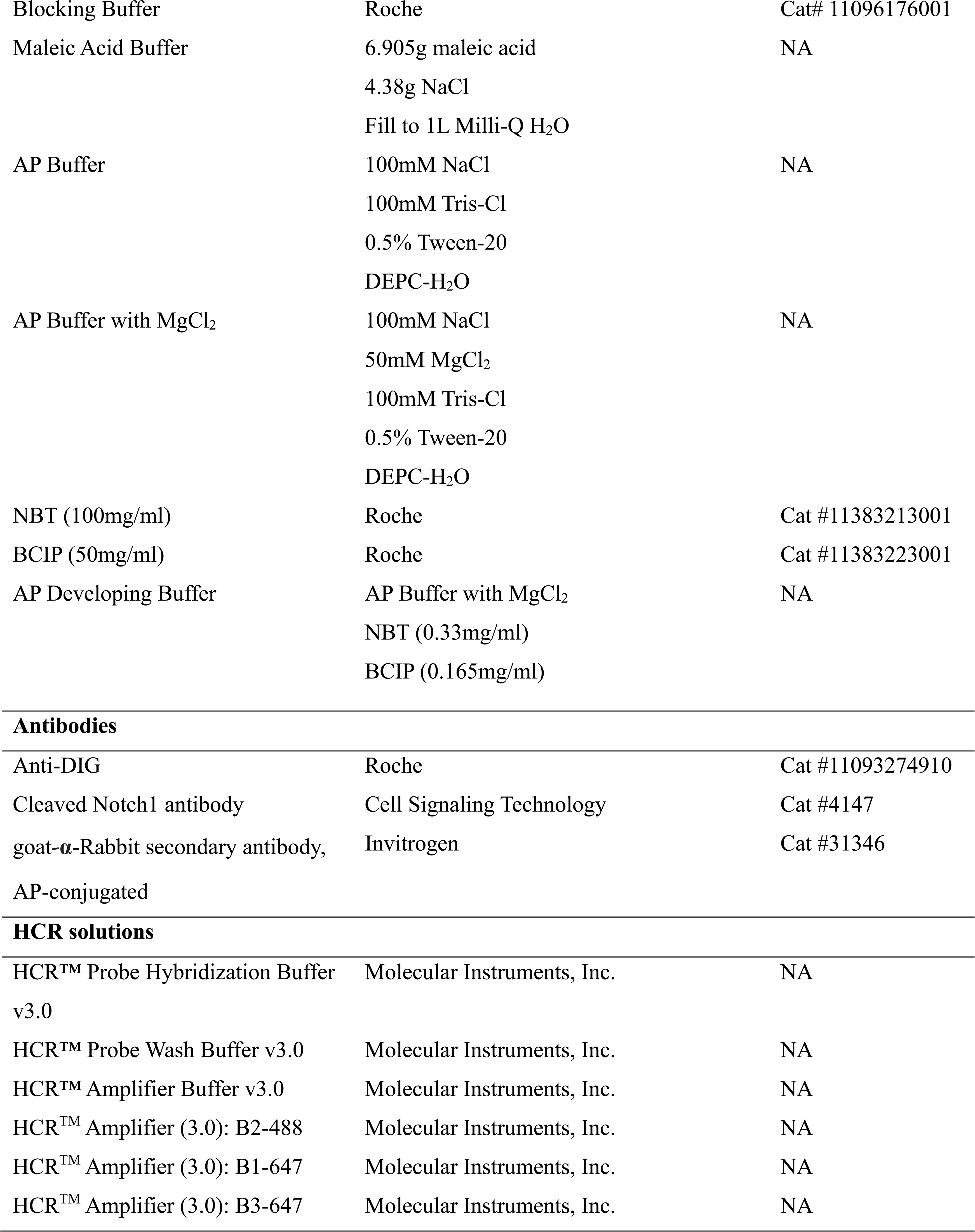
Key resources table

#### Animal collection and husbandry

Adult *Mnemiopsis leidyi* were collected in the Matanzas River estuarine system near St. Augustine and Beverly Beach, FL, and transported to the Whitney Laboratory. Animals were transitioned into 2/3X filtered sea water (FSW) (approximately 25 ppt) and kept in a self-circulating Kreisel system in constant light. Adults were fed several times per day with artemia brine shrimp enriched with S. Presso (Seclo^®^). Prior to spawning, adult animals were placed in the dark for 3 hours and then transitioned to glass finger bowls. When spawning began, roughly one hour later, embryos were collected in a 30-minute window and allowed to develop until 4 hours post fertilization (hpf), after which the vitelline membrane was manually removed with sharp forceps. Embryos were then allowed to grow up until 8hpf or 24hpf when they were fixed for *in situ* hybridization or HCR^TM^.

#### Gene Identification and Domain Analysis

*M. leidyi* genes were identified in the *Mnemiopsis* Genome Portal from gene identification numbers^18^. Orthology was confirmed via reciprocal BLAST against the NCBI database^49^ and PFAM analysis. PFAM architectures were evaluated using a combination of SMART^50^ and Prosite^51–53^. Domain significance was determined by E-value (SMART) or score threshold (Prosite) (see Supplementary File 1 for PFAM scores). Juxta-transmembrane domains were confirmed using TMHMM 2.0^54^ and the Conserved Domain Search (CDS) from NCBI^55–57^ (E-value ≤ 0.01). Nuclear localization signals (NLS) were identified with NL Stradamus^58^ (significant posterior probability ≥ 0.6). PEST domains were identified with EPestfind (http://emboss.bioinformatics.nl/cgi-bin/emboss/epestfind; threshold ≥ +5.0 score). Putative S1, S2, and S3 recognition sites were identified from previous publications^59^. *MlNotch* juxta-transmembrane domain (JTMD) was aligned with reported *TaNotch*, *NvNotch*, *HsNotch1, HsNotch4, HsNotch2, HsNotch3, DmNotch* JTMDs (see Supplementary File 2).

#### Phylogeny

Sequences for ctenophores (*Boroe ovata*, *Mnemiopsis leidyi*, and *Hormiphora Califoniensis*) and choanoflagellates (*Capsaspora owczarzaki* and *Mylnosiga fluctuans*) were added to existing alignments^3,16^ using MAFFT^60,61^ (see Supplementary File 3 for alignments). EGF repeat domains were excluded from the alignments due to their variability across species^3^. Bootstrapped maximum likelihood phylogenies were prepared with RAxMLGUI2.0 using a WAG substitution matrix and executing 500 ML tree inferences and thereafter a thorough ML search. All free model parameters were estimated by RAxML using a gamma model of rate heterogeneity. Bayesian analysis was performed using MrBayes 3.2.6 in Geneious^62^ using the WAG fixed model and gamma rate variation. Markov Chain Monte Carlo runs of 1,100,000 generations were calculated with trees sampled every 200 generations and with a prior burn-in of 110,000 generations. The best-scoring tree was visualized using FigTree (v1.4.2; http://tree.bio.ed.ac.uk/software/figtree) and annotated in Adobe Illustrator 2022. Juxtatransmembrane alignments were analyzed and visualized with Jalview^63^. AlphaFold predictions were made with ColabFold v1.5.5: AlphaFold2 using MMseqs2^64^ (see Supplementary File 3).

#### Developmental RNAseq for M. leidyi

Developmental RNA-seq data^21^ was accessed via the *Mnemiopsis* Genome Portal^18^ and plotted in Excel (see Supplementary File 5).

#### Cloning of in situ probe templates

Probe preparation and colorimetric *in situ* hybridization were conducted as previously described^65^, with slight modifications. Gene fragments were ordered with flanking SP7 and T7 promoters from Twist Biosciences, Inc. (see Supplementary File 6 for fragment sequences) and directly PCR amplified for probe synthesis using SP6 and T7 primers. Some genes were amplified from cDNA using gene specific primers. These were then ligated into Promega PGEM-T vector (A3600), then transformed and verified via colony PCR. Plasmids were purified using the ThermoFisher GeneJET Plasmid Miniprep kit (#K0503) and analyzed by Sanger sequencing. DIG-labled RNA probes were transcribed using Invitrogen MEGAscript SP6 (#AM1330) or T7 (#AM1334) kits according to manufacturer directions.

#### Colorimetric whole-mount in situ hybridization

Whole-mount *in situ* hybridizations were conducted as previously described^65^ with slight modifications. Animals were fixed in ACME solution^66^ for one hour at room temperature, then subsequently washed in PBS-Tween (see Supplementary File 7 for the detailed protocol). In brief, samples underwent a secondary fixation in 4% PFA and 0.8% glutaraldehyde for five minutes at room temperature, then transitioned into 4% PFA solution overnight at 4°C. The following day, samples were transitioned to 100% methanol and stored at -20°C. Samples were subjected to Proteinase K digestion and washes in glycine, triethanolamine, and acetic anhydride before proceeding with an overnight pre-hybridization at 63°C. Probes were hybridized using 1ng/µl concentrations at 63°C for 24–72 hours. Samples were transitioned into SSC and PBS-Tween solutions and then blocked in Roche Blocking Buffer (Roche, #11096176001) overnight at 4°C. The following day, samples were incubated with DIG-antibody (Sigma, #11093274910) diluted in Roche Blocking Buffer (Roche, #11096176001, 1:5000) overnight at 4°C. Samples were then washed in PBS-Tween and washed in alkaline phosphatase (AP) developing buffer (1M NaCl, 2M MgCl2, 1M Tris, Tween-20, NBT, BCIP) to develop DIG-labeled probes.

#### Hybridization chain reaction RNA fluorescence in situ hybridization (HCR RNA-FISH)

HCR probes were made using a customized Python code^67^ and ordered through IDT, Inc., and diluted to 100nM stock concentration. Animals were fixed at 8hpf using Rain-X® primary fixation^68^ for between 20–60 minutes at room temperature, transitioned into PBS-Tween and then subjected to an overnight secondary fixation in 4% PFA at 4°C. Samples were transitioned into 100% methanol and stored at -20°C until ready for use, after which they were transitioned back into PBS-Tween. We then followed a standard HCR^TM^ RNA-FISH protocol as described previously^23,69,70^, with slight modifications (see Supplementary File 8 for the detailed protocol). Samples were washed in HCR™ Probe Hybridization Buffer v3.0 (Molecular Instruments, Inc.) and incubated at 37°C. Probes were diluted to 20nM in HCR™ Probe Hybridization Buffer v3.0 (Molecular Instruments, Inc.) and were allowed to hybridize overnight at 37°C, then washed with HCR™ Probe Wash Buffer v3.0 (Molecular Instruments, Inc.). Samples were then washed in HCR™ Amplifier Buffer v3.0 (Molecular Instruments, Inc). Hairpins were heated at 95°C for 90 seconds, then allowed to cool to room temperature in the dark before being added to samples and allowed to amplify overnight at room temperature in the dark. Samples were then washed in 5X SSC and counterstained with Hoechst (1:1000).

#### Imaging

Colorimetric *in situ* samples were mounted in 80% glycerol and imaged on a Zeiss Imager M2. Images were post-processed in Adobe Photoshop 2022. HCR^TM^ samples were mounted in 5X SSC and imaged on a Zeiss 710 laser scanning confocal microscope. Confocal slices and Z-stacks were analyzed in FIJI^71^.

#### Pharmacological Inhibition

Embryos were de-membranated between 3–4hpf, then treated in 10µM DAPT (Sigma, D5942), 25µM LY411575 (Selleck Chemicals, S2714), and 1% DMSO (Sigma, D4540) in 2/3X FSW for approximately 4 hours. We then adapted a standard protocol for antibody staining of *Mnemiopsis leidyi* specimens (see Supplementary File 9 for the detailed antibody staining protocol). In brief, embryos were fixed in Rain-X® (200µl per 1ml) for 20 minutes, then washed into PBS-Tween and stored in 4% paraformaldehyde at 4°C overnight. The following day, embryos were washed 4 times in PBS-Triton, then permeabilized with 4 washes in PBT and blocked in 5% Normal Goat Serum (NGS) for 1 hour at room temperature. Cleaved Notch1 antibody (Cell Signaling Technology, #4147, 1:500) was diluted in 5% NGS and incubated with samples overnight at 4°C. Embryos were washed 7 times in PBT, then incubated with goat**-α**-Rabbit secondary antibody conjugated with AP (Invitrogen, #31346, 1:3500) and incubated overnight at 4°C. The specimens were then washed twice in AP developing buffer without MgCl_2_ (1M NaCl, 1M Tris, Tween-20), then washed twice in AP buffer with MgCl_2_ (1M NaCl, 2M MgCl_2_, 1M Tris, Tween-20), then developed in AP buffer with NBT and BCIP.

## Supplemental information

