## Supplementary material for "*Notch* expression during ctenophore development gives insight into its ancestral function": SupFile7_ISH_Mnemiopsis-leidyi.pdf

**Title:**

**Colorimetric whole-mount RNA in situ hybridization for the ctenophore *Mnemiopsis leidyi* at all life stages**

Adapted from Mitchell & Edgar et al. *Frontiers in Zoology* 2021 and Pang & Martindale *Cold Spring Harbor Protocols* 2008.

### Introduction:

Ctenophores, or comb jellies, are gelatinous marine invertebrates that have remarkable biological properties and hold a key phylogenetic position for understanding early animal evolution. Unfortunately, the delicate tissues of these animals complicate efforts to examine morphological and cellular features in fixed material. The bodies of ctenophores contain "mesoglea," an extracellular matrix filled with glycosaminoglycans which disrupt sample integrity in conventional aldehyde-, alcohol-, and acetone-based fixation protocols. Our previous fixation protocol was limited to small cydippid stage animals ($\lesssim$1 mm). The specialized fixation and secondary-fixation protocols described herein preserve morphology and allow for downstream molecular analyses such as immunohistochemistry and in situ hybridizations in animals ranging from 0.5 mm to at least 5 mm, but are not compatible with phalloidin labeling.

Here, we describe two protocols: 1) an improved fixation technique utilizing a commercially available silane-containing solution called Rain-X® and 2) a protocol for whole-mount RNA in situ hybridization (ISH) for the visualization of spatiotemporal gene expression downstream of this fixation. The secondary-fixation described in the first protocol is dispensable for some downstream applications (such as immunohistochemistry) not detailed here; however, this secondary-fixation is essential for harsher treatments such as whole-mount in situ hybridization, as shown in the second protocol.

**BASIC PROTOCOL 1: Ctenophore Fixations**

Small adult ctenophores of many species are called "cydippids," and in some species cydippids continue to grow differentially into a morphologically modified adult lobate stage, which may present added difficulties in morphological preservation during fixation. Adult tissue samples are first fixed with Rain-X®, and if necessary are then secondary-fixed with 4% paraformaldehyde (PFA) and 0.2% glutaraldehyde (GTA) to improve tissue durability. Following fixation, samples are washed in PBS-Tween (PTw) and phosphate-buffered saline (PBS), after which they are dehydrated and stored in 100% methanol (MeOH) at temperatures between 4ºC and -20ºC. Note that tissues remain fragile throughout these procedures and special care must be taken during pipetting and exchanging solutions to ensure sample preservation.

| **Materials** | **Catalog Number** | **Recipe** | |
|---|---|---|---|
| Full-strength filtered sea water (FSW) | NA | —— | |
| Rain-X® | NA | —— | |
| 25% Glutaraldehyde (GTA)<br>Electron Microscopy Sciences | #16216-10 | —— | |
| 16% Paraformaldehyde (PFA)<br>Electron Microscopy Sciences | #15710 | —— | |
| Milli-Q Water (MQ-$H_2O$) | NA | —— | |
| Diethyl pyrocarbonate (DEPC) | #D5758 | —— | |
| 10X Phosphate Buffer Solution (PBS)[1,2] | NA | *For 1L:*<br>2.23g $NaH_2PO4$<br>10.60g $Na_2HPO4$<br>102.2g NaCl<br>1L MQ-$H_2O$ | *Final concentration:*<br>18.6mM $NaH_2PO4$<br>84.1mM $Na_2HPO4$<br>1,750mM NaCl<br>—— |
| 1X Phosphate Buffer Solution (PBS) | NA | *For 50ml:*<br>5ml 10x PBS<br>45ml MQ-$H_2O$ | *Final concentration:*<br>1X<br>—— |

| | | | |
|---|---|---|---|
| Tween<br>Sigma | #P2287 | —— | |
| 1X PBS-Tween (PTw)[1] | NA | *For 1L:*<br>100ml 10x PBS<br>900ml MQ-$H_2O$<br>1ml Tween | |
| 100% Methanol (MeOH)<br>Fisher | #A412-4 | —— | |
| 60% MeOH | NA | *For 10ml:*<br>6ml MeOH<br>4ml MQ-$H_2O$ | |
| 80% MeOH | NA | *For 10ml:*<br>8ml MeOH<br>2ml MQ-$H_2O$ | |
| Ctenophore Secondary-fixation Solution 1[3] | NA | *For 5ml:*<br>40μl 25% GTA<br>1.25ml 16% PFA<br>3.35ml 1X PBS | *Final concentration:*<br>0.2% GTA<br>4.0% PFA<br>—— |
| Ctenophore Secondary-fixation Solution 2[3] | NA | *For 5ml:*<br>1.25ml 16% PFA<br>3.75ml 1X PBS | *Final concentration:*<br>4.0% PFA<br>—— |

Table 1.

[1] DEPC treatment (RNAse-free):

1. For 1L solutions, add 1mL of DEPC to an autoclave-save bottle under a fume hood with a 1mL pipette with unfiltered tips, popping the tip off into the bottle after administering the DEPC
2. Close the bottle tightly and wrap parafilm around lid to ensure leakage doesn't occur
3. Shake the bottle vigorously under the hood for 30 seconds
4. Remove parafilm and incubate for 1 hour
5. Loosen lid and autoclave for 45 mins

[2] PBS (10x): Instructions for making solution

6. Mix phosphates in 800mL of MQ-$H_2O$ in an autoclave-safe bottle
7. Check pH – reading should be 7.4 +- 0.4

8. Adjust pH with NaOH or HCl if necessary
9. Add NaCl
10. DEPC treat and autoclave for 45 mins

[3] Store solutions at 4ºC. They will keep for 1–2 days.

| Materials | Catalog Number |
|---|---|
| 6-well plate/24-well plate<br>Falcon | #353046 |
| 50ml conical screw-cap tubes<br>Thermo | #339652 |
| 5ml conical screw-cap tubes<br>Mtcbiotech | #C2540 |
| 1.5ml microcentrifuge tubes<br>Fisher | #05-408-129 |
| Syringe<br>BD | #309659 |
| 22-gauge needle<br>BD | #305111 |
| Parafilm | —— |
| Ice | —— |

Table 2.

**Primary Fixation**

1. Transfer live animal sample(s) in a known volume of full-strength filtered sea water (FSW).

   *Use the total volume of FSW + animal(s) as the "sample volume" when calculating downstream proportional volumes.*

   *We have successfully used 4-, 6-, 12-, and 24-well plates as well as 50ml conical tubes.*

   *When transferring animals and adding solutions, use a dissecting scope to ensure that animals are not inadvertently removed.*

*At each step in this protocol, make sure to leave enough liquid in each well so that samples do not dry out between washes.*

2. For animals smaller than 5 mm, add 200µl Rain-X® per ml sample volume (i.e. final volume with FSW + animals + Rain-X® = 1.2 ml).

   *Make sure samples are completely covered by liquid to avoid drying out. Mix gently but thoroughly by pipetting only the liquid—this is important to prevent animals from sticking together. Avoid overcrowding samples in wells or tubes.*

   *For larger-format tubes (i.e. 50 ml conical tubes), gentle inversion works well.*

3. Fix sample(s) in Rain-X® overnight at 4ºC (preferred).

   *Samples can also be fixed for 1–2 hours at room temperature (RT). Place samples on a gentle rocker during fixation. To prevent samples from clumping together and maintain complete submersion in the RainX®/FSW solution during this initial fixation, gently pipette the liquid up and down in the wells approximately every 15 minutes. Do not pipette animals.*

4. Remove as much of the Rain-X® mixture as possible without removing sample(s).

   *This is easier to do under a dissecting microscope.*

5. Wash 1 time with 1X PBS.

   *At this point, samples may be washed several times with PBS and used directly for staining and imaging, immunohistochemistry, etc. Samples intended for such uses may be stored for several days in PTw at 4ºC. The subsequent steps are indispensable for RNA in situ hybridizations.*

   *For larger-format tubes (i.e. 50 ml conical tubes), transferring the samples to a fresh tube with a widened transfer pipette at this point can be useful for removing Rain-X® that may coat the sides and lid of the tube.*

6. Remove as much 1X PBS as possible without removing sample(s).
7. Wash 2 times in PTw for 5 minutes each.
8. Remove as much PTw as possible without removing sample(s).

**Prepare secondary-fixation solutions**

1. Withdraw GTA stock solution from its container using a needle on a syringe to pierce the vial stopper and transfer solution to a plastic microfuge tube.

   *We have found GTA to be indispensable for preservation of morphology during ISH. GTA is hazardous, so use caution during this step. Pipette GTA in a fume hood and dispose solids and liquids in designated hazardous waste containers.*

2. Make Ctenophore secondary-fixation Solution 1 and Ctenophore secondary-fixation Solution 2.

**Secondary-fixation**

1. Add Ctenophore Secondary-Fixation Solution 1 to your sample(s) and gently mix. Fix for 5 minutes at RT.

   *Be sure to add enough solution to completely cover samples.*

   *Alternatively, you can leave animals in Ctenophore Secondary-Fixation Solution 1 overnight at* 4ºC.

2. Remove as much of the Ctenophore Secondary-Fixation Solution 1 as possible without removing sample(s).

   *Discard in GTA waste.*

3. Add Ctenophore Secondary-Fixation Solution 2 and gently mix. Fix 1 hour at RT.

   *Be sure to add enough solution to cover samples.*

   *Alternatively, you can leave animals in Ctenophore Secondary-Fixation Solution 1 overnight at* 4ºC and skip *Ctenophore Secondary-Fixation Solution 2.*

4. Wash 2 times in PTw for 5 minutes each.
5. Wash 2 times in PBS for 5 minutes each.
6. Wash 3 times with Milli-Q water for 5 minutes each.
7. Wash 2 times with RNAse-free 50% MeOH for 5 minutes each.
8. Wash 1 time with RNAse-free 80% MeOH for 5 minutes each.
9. Wash at least 2 times with RNAse-free 100% MeOH for 5 minutes each, then move to a fresh container of 100% MeOH.
10. Store sample(s) at -20ºC.

*After 1 hour, samples may be used for in situ hybridizations or stored for several months at -20ºC until ready for use. If using a loose-lidded container such as a multiwell-plate, make sure the container is closed and well-sealed with parafilm, otherwise MeOH will evaporate in the freezer over time.*

**BASIC PROTOCOL 2: Ctenophore whole-mount in situ hybridization**

Here, we describe multi-day pretreatment and prehybridization steps necessary for labeled riboprobes to hybridize to RNA and undergo a colorimetric alkaline phosphatase (AP) reaction for detecting spatiotemporal gene expression. Pretreatment steps consist of transitioning samples from MeOH to PTw and then permeabilizing tissue using Proteinase-K digestion. (NB: Proteinase-K digestion is incompatible with performing immunohistochemistry as a secondary label on the same samples as it may damage target antigens.) Once the digestion is stopped, samples are refixed in 4% PFA and then washed in PTw. After incubating samples in hybridization buffer, probes are added and allowed to hybridize. Probe hybridization is detected with a colorimetric reaction catalyzed by an AP-conjugated anti-digoxigenin antibody. Antisense riboprobes should be prepared ahead of time; we do not provide a protocol for design or synthesis of these probes. The gold-standard negative control for ISH is a labeled sense (rather than antisense) probe for the gene of interest. It is best to include 1–2 known positive control probes on a plate when evaluating any new probe.

Day 1

| Materials | Catalog Number | Recipe |
|---|---|---|
| 100% MeOH | See Table 1. | —— |
| 60% MeOH | NA | *For 10ml:*<br>6ml 100% MeOH<br>4ml PTw |
| 30% MeOH | NA | *For 10ml:*<br>3ml 100% MeOH<br>7ml PTw |
| Proteinase-K(protk) [4]<br>Invitrogen | #25530049 | 20mg/ml |

| | | | |
|---|---|---|---|
| PTw-Proteinase-K | NA | *For 10ml:*<br>5µl 20mg/ml protk<br>Fill to 10ml PTw | *Final concentration:*<br>0.01mg/ml protk<br>—— |
| RNAse-free glycine<br>Sigma | #410225 | —— | |
| PTw-glycine[5] | NA | *For 50ml:*<br>0.1g glycine<br>50ml PTw | *Final concentration:*<br>2mg/ml glycine |
| Triethanolamine (TEA)<br>Sigma | #90279 | —— | |
| PTw-TEA | NA | *For 10ml:*<br>100µl TEA<br>Fill to 10ml | *Final concentration:*<br>1% (10µl/ml) TEA |
| Acetic anhydride<br>Sigma | #320102 | —— | |
| 16% PFA | See Table 1. | —— | |
| PTw-PFA | NA | *For 5ml:*<br>1.25ml 16% PFA<br>3.35ml PTw | *Final concentration:*<br>4% PFA<br>—— |
| Salmon Sperm DNA(SSD) (10mg/ml)<br>Invitrogen | #AM9680 | —— | |
| Formamide<br>Sigma | #FX0420 | —— | |
| Tween | #P2287 | —— | |
| Heparin(20mg/mL) | NA | —— | |

| | | | |
|---|---|---|---|
| SDS | #BP166-500 | —— | |
| 20% SDS solution[6] | NA | *For 1L:*<br>100g SDS<br>500mL MQ-$H_2O$ | *Final concentration:*<br>20% SDS |
| SSC(20X) pH 4.5[7] | NA | *For 1L:*<br>175.3g NaCl<br>88.2g Na citrate<br>1L MQ-$H_2O$ | *Final concentration:*<br>3 M NaCl<br>0.3 M Na citrate |
| DEPC $H_2O$[1] | NA | *For 1L:*<br>1mL DEPC<br>Fill to 1L MQ-$H_2O$ | *Final concentration:*<br>0.1% DEPC |
| In situ hybridization buffer with salmon sperm DNA (ISHB + SSD) | NA | *For 40ml:*<br>20mL Formamide<br>100µl Heparin<br>200µl SSD<br>2mL SDS<br>10mL SSC (pH 4.5)<br>200µl Tween<br>Fill to 40mL DEPC $H_2O$ | *Final concentration:*<br>50% Formamide<br>50µg/mL Heparin<br>100µg/mL SSD<br>1% SDS<br>5X SSC (pH 4.5)<br>0.1% Tween |

**Table 3.**

[4]Proteinase-K should be kept on ice when not in use.

[5]Aliquots of 2mg/ml glycine in PTw can be kept at -20 ºC

[6]Add 100g SDS to 400mL MQ-$H_2O$, head to 68 ºC and mix with a stir bar until dissolved. Fill to 500mL with MQ-$H_2O$. Use a face mask to precent inhalation. DO NOT AUTOCLAVE

[7]Add NaCl and Na Citrate to 800ml MQ-$H_2O$ in autoclave safe bottle, pH adjust to 4.5 with acid and/or base, fill to 1L with more MQ-$H_2O$ and autoclave for 45 mins

| Materials | Catalog Number |
|---|---|
| 6-well plate/24-well plate | See Table 2. |
| 1.5ml microcentrifuge tubes | See Table 2. |
| Pipette with cut tip | See Table 2. |
| Ice | NA |

Table 4.

**Pretreatment**

1. Remove stored sample(s) from -20ºC and carefully transfer fixed samples to new wells with 100% MeOH.

   *500 µl 100% MeOH in a 24-well plate is sufficient for 50-200 cydippids or up to 10 small lobates. All subsequent washes described herein are 500 µl unless otherwise specified.*

2. Remove as much 100% MeOH as possible without exposing sample(s) to air, then add 500–800µl 60% MeOH for 5 minutes.
3. Remove as much 60% MeOH as possible without exposing sample(s) to air, then rehydrate with 500–800µl 30% MeOH for 5 minutes.
4. Remove as much 30% MeOH as possible without exposing sample(s) to air.
5. Wash 5 times with 500–800µl PTw for 5 minutes each.

**Proteinase-K Digestion**

1. Digest sample(s) with 500µl PTw-Proteinase-K at RT.

   *Digest for 5 minutes for cydippids, small lobates, and adult tissues. Increase digestion to 10 minutes for embryos. Optimization of this step may be required for other Proteinase-K sources or special sample types.*

2. Remove Proteinase-K, then wash 2 times with PTw-glycine solution for 5 minutes each.

3. Remove PTw-glycine solution, then wash 2 times with 500µl PTw-TEA for 5 minutes each. Leave the specimens in the PTw-TEA after the second wash
4. Add 1.5µl acetic anhydride and mix by carefully swirling the plate or on a rocker for 5 minutes. The acetic anhydride will appear as a small bead at the bottom of the well
5. After 5 minutes has passed, add another 1.5µl of acetic anhydride and swirl for another 5 minutes.

   *Steps involving concentrated acetic anhydride should be performed under a hood.*
6. Remove the previous solution, then wash 2 times with fresh PTw for 5 minutes each.

**Post-Fixation**

1. Remove PTw and postfix samples in PTw-PFA for 1 hour at RT on a rocker with a gentle cycle.
2. Remove PTw-PFA and wash 5 times in PTw for 5 minutes each.

**Prehybridization**

1. Make in situ hybridization buffer with salmon sperm DNA (ISHB + SSD).

   *ISHB may be stored at -20℃ for several months.*
2. Wash sample(s) 2 times with 500 µl ISHB + SSD for 5 minutes each.

   *For all steps prior to introducing the riboprobes, use ISHB with salmon sperm DNA.*
3. Place sample(s) in hybridization oven at 63ºC for at least 1 hour.

   *This step is flexible—samples can prehybridize for 1–24 hours. To prevent evaporation, place samples in a secondary container with a damp paper towel to humidify the chamber.*

Day 2

| Materials | Catalog Number | Recipe | |
|---|---|---|---|
| In situ hybridization buffer without salmon sperm DNA (ISHB - SSD) | NA | *For 40ml:*<br>20mL Formamide<br>100µl Heparin<br>2mL SDS<br>10mL SSC(pH 4.5)<br>200µl Tween | *Final concentration:*<br>50% Formamide<br>50µg/mL Heparin<br>1% SDS<br>5X SSC(pH 4.5)<br>0.1% Tween |

| | | |
|---|---|---|
| | | Fill to 40mL DEPC $H_2O$ |
| Riboprobe stocks | NA | —— |

Table 5.

| Materials | Catalog Number |
|---|---|
| 50ml conical screw-cap tube | See Table 2. |
| 6-well or 24-well plate containing sample(s) | See Table 2. |

Table 6.

**Hybridization**

1. Dilute riboprobe stocks to a final concentration of 1ng/µl in ISHB + SSD in a centrifuge tube.
2. Denature probes in a heat block at 80–90ºC for 10 minutes.
3. Take the sample plate from the hybridization oven and remove hybridization buffer from each well containing samples, taking care not to expose the samples to air.

   *Discard in formamide waste.*
4. Add 500µl of denatured probe solution to each well.
5. Return the plate to the humidified chamber in the oven at 63ºC and allow to hybridize.

   *We recommend hybridizing for a minimum of 16 hours. Typically we hybridize for 24–48 hours, and have hybridized up to 3 days without disrupting quality.*

Day 3

| **Materials*** | **Catalog Number** | **Recipe** |
|---|---|---|
| ISHB - SSD | NA | See Table 5. |
| 2X saline sodium-citrate buffer (SSC), pH 7<br>Sigma | #S6639 | —— |

| | | |
|---|---|---|
| ISHB : 2X SSC (75 : 25) | NA | *For 40ml:*<br>30ml ISHB - SSD<br>10ml 2X SSC |
| ISHB : 2X SSC (50 : 50) | NA | *For 40ml:*<br>20ml ISHB - SSD<br>20ml 2X SSC |
| ISHB : 2X SSC (25 : 75) | NA | *For 40ml:*<br>10ml ISHB - SSD<br>30ml 2X SSC |
| 0.05X SSC, pH 7 | NA | *For 50ml:*<br>1.25 ml 2X SSC (pH 7)<br>Fill to 50ml MQ-$H_2O$ |
| PTw | NA | See Table 1. |
| Blocking Buffer(10X)<br>Roche | #11096176001 | |
| 1X Blocking buffer[9] | NA | *For 10mls:*<br>1ml 10X Roche Blocking Buffer<br>Fill to 10mls with Maleic Acid Buffer |
| Maleic Acid Buffer[10] | NA | *For 1L:*<br>6.905g Maleic acid<br>4.38g NaCl<br>Fill to 1L MQ-$H_2O$ |
| AP-conjugated anti-digoxigenin<br>Roche | #11093274910 | —— |

Table 7.

**8** It is best to prepare these solutions at least the day before. ISHB can be stored at -20ºC. SSC can be stored at RT, however if mixed with ISHB it should be stored at 20ºC.

**9** 10X Roche block can be stored at 4 ºC but 1X in Maleic acid should be made fresh before blocking step.

**10** Add Maleic acid and NaCl to 400mLs of MQ-$H_2O$, adjust pH to 6.5 with NaOH pellets, adjust to 7.4 with 5M NaOH, adjust to 7.5 with 1M NaOH and fill to 1L with MQ-$H_2O$ and autoclave.

| **Materials** | **Catalog Number** |
|---|---|
| 50ml conical screw-cap tubes | See Table 2. |

| | |
|---|---|
| 6-well or 24-well plate containing sample(s) | See Table 2. |

Table 8.

**Hybridization (continued)**

1. Heat the following wash solutions to 60ºC in the hybridization oven: ISHB, all listed mixtures of ISHB : SSC, 2.0X SSC, and 0.05X SSC.
2. Remove probes by pipetting solution of wells.
   *Used probes can be saved and stored at -20ºC for re-use.*
3. Wash samples in 500µl 100% ISHB for 20 minutes in the hybridization oven at 60ºC.
   *RNase-free conditions are unnecessary beyond this point. Salmon sperm DNA is not required in ISHB for this or any subsequent step. The humidified chamber is also no longer necessary.*
4. Remove 100% ISHB, then wash with 500µl ISHB : 2X SSC (75 : 25) in hybridization oven at 60ºC for 20 minutes.
5. Remove ISHB : 2X SSC (75 : 25), then wash with 500µl ISHB : 2X SSC (50 : 50) in hybridization oven at 60ºC for 20 minutes.
6. Remove ISHB : 2X SSC (50 : 50), then wash with 500µl ISHB : 2X SSC (25 : 75) in hybridization oven at 60ºC for 20 minutes.
7. Remove ISHB : 2X SSC (25 : 75), then wash 3 times with 500µl 2X SSC in hybridization oven at 60ºC for 20 minutes each.
8. Remove 2X SSC, then wash 3 times in 0.05X SSC in hybridization oven at 60ºC for 20 minutes.
   *The concentration of SSC and length of washes are essential for stringency of probe hybridization.*
9. Wash samples in PTw 5 times at RT for 5 minutes each.
   *This step is flexible; once samples are in PTw, they may be washed longer than 5 minutes.*

**Detection for chromogenic ISH using AP-conjugated antibody**

1. Add 500µl of blocking buffer and incubate at 4ºC overnight.

   *Alternatively, blocking buffer incubation may be done for 1 hour at RT.*

2. Dilute AP-conjugated anti-digoxigenin antibody 1 : 5000 in blocking buffer solution.

3. Add 500µl of diluted antibody to each sample.

4. Incubate samples overnight at 4ºC.

Day 4

| **Materials** | **Catalog Number** | **Recipe** | |
|---|---|---|---|
| PTw | NA | See Table 1. | |
| 1X PBS | NA | See Table 1. | |
| 1M $MgCl_2$ | NA | | |
| 5M NaCl | NA | | |
| 1M Tris-Cl(pH 9.5) | NA | | |
| AP buffer **without** $MgCl_2$ [11] | NA | *For 10ml:*<br>200µl NaCl(5M)<br>1mLTris-Cl(1M)<br>250µl Tween<br>Fill to 10ml DEPC $H_2O$ | *Final concentration:*<br>100mM NaCl<br>100mM Tris-Cl<br>0.5% Tween |
| AP buffer **with** $MgCl_2$[11] | NA | *For 10ml:*<br>500µl $MgCl_2$(1M)<br>200µl NaCl(5M)<br>1mLTris-Cl(1M)<br>250µl Tween<br>Fill to 10ml DEPC $H_2O$ | *Final concentration:*<br>50mM $MgCl_2$<br>100mM NaCl<br>100mM Tris-Cl<br>0.5% Tween |
| DEPC $H_2O$[1] | NA | See Table 3. | |

| | | | |
|---|---|---|---|
| NBT(100mg/ml)<br>Roche | #11383213001 | —— | |
| BCIP(50mg/ml)<br>Roche | #11383221001 | —— | |
| AP developing buffer[12] | | *For 10ml:*<br>33μl BCIP<br>66μl NBT<br>Fill to 10ml with AP with $MgCl_2$ | Final concentration<br>0.165 mg/ml<br>0.33 mg/ml |
| EDTA(0.5M)[13] | #E5134 | *For 1L:*<br>186.1 g EDTA<br>Fill to 1L MQ-$H_2O$ | |
| Glycerol<br>Fisher | #BP229-1 | | |

Table 9.

[11] Needs to be made fresh before use

[12] The molecular weight of NBT is 2X that of BCIP so at this concentration you are using 8 molecules of NBT per 1 molecule of BCIP (final reaction product occurs faster)

[13] A recommended pH of this solution is 8.0, so NaOH may need to be used. This solution can be autoclaved, but it is not necessary for this step of the protocol. Dilute to 50mM in MQ-$H_2O$ before use in the protocol to stop the reaction .

| Materials | Catalog Number |
|---|---|
| 6-well or 24-well plate containing sample(s) | See Table 2. |

Table 10.

**Detection (continued)**

1. Wash sample(s) 12 times with 500μl PTw at RT for 10 minutes each.
2. Wash sample(s) 3 times with 500μl PBS at RT for 10 minutes each.

3. Wash sample(s) 2 times with 500µl AP developing buffer **without** $MgCl_2$ at RT for 5 minutes each.
4. Wash sample(s) 2 times (2X) with 500µl AP developing buffer **with** $MgCl_2$ at RT for 5 minutes each.
5. Remove and replace with 500µl fresh AP developing buffer, then cover in foil or place in the dark for the reaction to occur at RT.

   *Plates may be placed on a gentle rocker.*
6. Monitor the reaction every 30 minutes for the first day of development. A dark blue-purple precipitate will form in cells that express the gene of interest.

   *Development can take as little as 30 minutes or as long as a few days; this timing will be probe-specific and must be determined empirically and monitored closely. The AP developing buffer should start off slightly yellowish and turn purple over time. Replace AP developing buffer before it turns purple.*

**Stopping AP reaction**

1. When sample(s) have been stained sufficiently, stop the reaction by removing AP developing buffer and washing 3 times with 500µl of 50mM EDTA for a minimum of 20 minutes each wash.
2. Remove EDTA and wash sample(s) 3 times with 500µl PTw for 5 minutes each.
3. Remove PTw and wash sample(s) 3 times with 500µl 1X PBS for 5 minutes each.
4. Mount sample(s) in 40–80% glycerol diluted in PBS and image.
