## Supplementary material for "*Notch* expression during ctenophore development gives insight into its ancestral function": SupFile8_Ctenophore_HCR_protocol.docx

**Hybridization Chain Reaction RNA fluorescence *in situ* hybridization (HCR RNA-FISH) in ctenophores**

version 2: written by Cezar Borba

Day 1: Ctenophore Fixations

| **Solutions** | **Recipe** | **Sample** |
| --- | --- | --- |
| Full-strength filtered sea water (FSW) | NA | NA |
| Rain-X® | NA | NA |
| 16% Paraformaldehyde (PFA) | NA | NA |
| 1X Phosphate Buffer Solution-Tween (PBST) | 1X Phosphate Buffer Solution (PBS)  0.1% Tween  H_2_O | For 50 mL:  5 mL 10X PBS  50 μL Tween  ~ 45 mL H_2_O |
| 100% Methanol (MeOH) | NA | NA |
| Ctenophore Post-fixation Solution | 4% PFA  PBST | For 2 mL:  500 μL 16% PFA  1.5 mL PBST |

Primary Fixation

1. Transfer live animal sample(s) to a known volume of FSW.
2. Add 200 μL Rain-X® per ml total volume FSW and animals. Fix for 1 hour at room temperature (RT)

*I usually have them in 3 mL in a 6-well plate for the quantity, so I add 600 μL*

3. Remove 25% of Rain-X® mixture and replace with PBST. Incubate for 5 minutes at RT.

*Since I would have 3600 μL at this point, I would replace 900 μL*

4. Remove 33% of Rain-X® mixture and replace with PBST. Incubate for 5 minutes at RT.

*Continuing, I would replace 1200 μL*

5. Remove 50% of Rain-X® mixture and replace with PBST. Incubate for 5 minutes at RT.

*Continuing, I would replace 1800 μL*

6. Do three 5-minute washes with PBST at RT

*Remove as much solution as possible without removing samples, add PBST (~1 mL in 24-well, ~2mL in 6-well). Since I do this in 6-well plates, I tend to leave 1 mL, then add 2 mL of PBST. Then, I just replace 2 mL with new PBST two times.*

Post-fixation

1. Add Ctenophore Post-fixation Solution to your sample(s) and gently mix. Fix for 45-60 minutes at RT.

*I just leave 1500 μL of PBST and directly add 500 μL of 16% PFA while gently mixing. Otherwise, I create a solution of 8% PFA in PBST and replace half the well solution with it.*

2. Do three 5-minute washes with PBST at RT.

*Same as step 7 in primary fixation. Sometimes, during the MeOH part (next steps), a white precipitant appears; to avoid that, move the animals to a fresh well with PBST using as little of the previous volume as possible.*

3. Remove 25% of PBST and replace with MeOH. Incubate for 5 minutes at RT.
4. Remove 33% of PBST and replace with MeOH. Incubate for 5 minutes at RT.
5. Remove 50% of PBST and replace with MeOH. Incubate for 5 minutes at RT.
6. Do at least two 5-minute washes with MeOH at RT.

*Washes done similar to step 7 in primary fixation. I tend to do 2 or 3 washes.*

7. Store sample(s) at -20ºC overnight.

Day 2: Hybridization step

| **Solutions** | **Recipe** | **Sample** |
| --- | --- | --- |
| 100% MeOH |  |  |
| 1X Phosphate Buffer Solution-Tween (PBST) | 1X Phosphate Buffer Solution (PBS)  0.1% Tween  H_2_O | For 50 mL:  5 mL 10X PBS  50 μL Tween  ~ 45 mL H_2_O |
| Probe Hybridization Buffer (PHB) | Purchased from molecular instruments |  |

1. Put an aliquot of PHB in 37ºC incubator.
2. Remove stored sample(s) from -20ºC and carefully transfer desired amount of fixed animals to new wells with 100% MeOH at RT.

*With 24hr cydippids, I’ve done anywhere from 15-200, depending how many I have to use*

3. Remove 25% of well solution and replace with PBST. Incubate for 5 minutes at RT.
4. Remove 33% of well solution and replace with PBST. Incubate for 5 minutes at RT.
5. Remove 50% of well solution and replace with PBST. Incubate for 5 minutes at RT.
6. Do three 5-minute washes with PBST at RT.
7. Remove 50% of well solution and replace with pre-heated PHB. Incubate for 15 minutes at 37ºC
8. Remove as much of well solution as possible without drying out sample(s) and replace with pre-heated PHB. Incubate for 30 minutes at 37ºC.
9. While waiting for step 8 to finish, create probe solution by adding the probe(s) to pre-heated PHB in separate tube/well (2 μL of each probe/100 μL of PHB; 20 nM probe solution). Incubate at 37ºC.

*Since I used 24-hr cydippids, I’ve been just prepping these in small tubes at 50-100 μL.*

10. Remove as much of well solution as possible without drying out sample(s) and replace with probe solution. Incubate overnight at 37ºC.

*I’ve been pipetting up the animals in the smallest volume possible (usually 8-10 μL*) *and putting them in the small tubes with the probe solution.*

Day 3: Amplification step

| **Solutions** | **Recipe** | **Sample** |
| --- | --- | --- |
| Probe Wash Buffer (PWB) | Purchased from molecular instruments |  |
| Amplification Buffer | Purchased from molecular instruments |  |
| 20x sodium chloride sodium citrate (20xSSC) | 3 M NaCl  0.3 M sodium citrate | For 50 mL:  8.77 g NaCl  4.41 g sodium citrate  Fill up to 50 mL with H_2_O |
| 5x sodium chloride sodium citrate-tween (5xSSCT) | 5x SSC  0.1% Tween | For 40 mL:  10 mL 20xSSC  40 μL Tween  ~30 mL H_2_O |

Probe Washing

1. Put an aliquot of PWB in 37ºC incubator, an aliquot of amplification buffer at RT, and pre-set a heat block for 95ºC.
2. Remove probe solution from samples (and if possible, save in -20ºC; can be re-used) and replace with pre-heated PWB. Incubate for 15 minutes at 37ºC.
3. Do three 15-minute washes with pre-heated PWB at 37ºC.
4. Remove 25% of PWB and replace with 5xSSCT. Incubate for 5 minutes at RT.
5. Remove 33% of PWB and replace with 5xSSCT. Incubate for 5 minutes at RT.
6. Remove 50% of PWB and replace with 5xSSCT. Incubate for 5 minutes at RT.
7. Do one-to-three 5-minute washes with 5xSSCT at RT.
8. Remove as much of well solution as possible without drying out sample(s) and replace with amplification buffer. Incubate at RT.

Hairpin preparation (hairpins should be kept on ice and protected from light while being prepared)

1. Pipette h1 and h2 of each hairpin into individual tubes (2 μL/100 μL final volume)
2. Place each hairpin tube on 95ºC heat block for 90 seconds.
3. Incubate hairpin tubes at RT in the dark for 30 minutes.
4. Create hairpin solution by adding hairpins to amplification buffer.

*Since I use 24hr cydippids, I’ve been prepping these in small tubes at 100 μL, sometimes even 50 μL*

5. Remove as much of well solution as possible without drying out sample(s) and replace with hairpin solution. Incubate at RT overnight in the dark.

*I’ve been pipetting up the animals in the smallest volume possible (usually 8-10 μL) and putting them in the small tubes with the hairpin solution.*

Day 4: Imaging

| **Solutions** | **Recipe** | **Sample** |
| --- | --- | --- |
| 20x sodium chloride sodium citrate (20xSSC) | 3 M NaCl  0.3 M sodium citrate | For 50 mL:  8.77 g NaCl  4.41 g sodium citrate  Fill up to 50 mL with H_2_O |
| 5x sodium chloride sodium citrate-tween (5xSSCT) | 5x SSC  0.1% Tween | For 40 mL:  10 mL 20xSSC  40 μL Tween  ~30 mL H_2_O |
| Counter stain solution*  *This is the solution I use, edit as seen fit. If not counter staining, just use 5xSSCT. | 5xSSCT  2.86 μM DAPI  2x Cell Mask | For 7 mL:  2 μL of 10 mM DAPI  14 μL of 1000x Cell Mask  ~7 mL of 5xSSCT |

1. Remove as much of the hairpin solution as possible without drying out sample(s) (if possible, save in -20ºC; can be re-used, although I haven’t tested this myself) and replace with 5xSSCT. Incubate for 5 minutes at RT in dark.
2. Do a 5-minute wash with 5xSSCT at RT in dark.
3. Remove half the well solution and replace it with counter stain solution. Incubate at RT in dark.
4. Do a 30-minute wash with 5xSSCT at RT in dark.
5. Do a 5-minute wash with 5xSSCT at RT in dark, then it’s ready to mounted and imaged.
