## Supplementary material for "*Notch* expression during ctenophore development gives insight into its ancestral function": SupFile9_Mnemiopsis-Antibody-Protocol.pdf

### Mnemiopsis Cydippid Antibody Staining Protocol

**updated 1/19/22 -  copy for Brent and Will

##### Materials:

- Using 1-3 mm cydippids
- RainX
- 3 spot glass dish — marked as fixative safe OR 24 well plate ( something with wells under 5mLs)
- PBT
- PBS 1x
- Pipette tips
- Scissors - to cut pipette tips if using large animals
- 0.2% PTx
- 5% NGS in PBT

##### Procedure :

1. Prepare and “RAIN Fix" Cydippids
    1. Select cydippids over 1mm for fixation
    2. Place cydippids in well in 3 spot glass dish in 1000µl UV Filtered Full Strength Sea Water (1x)
    3. Administer 200µl of RainX to the well
        1. Pay careful attention that the RainX is homogenized so that all components of the solution can reach the tissue
    4. Cover the dish and leave the animal to fix for 1 hour @ RT
    5. Wash fix out 3X for 10 minutes with 1X PBS
2. Permeabilization ( i ) : 500-1000µl washes
    1. Drain 1x PBS as much as possible out of well from previous step
    2. Add 0.2% Triton in PBS (PTx)
    3. Let Sit for 10 mins, then remove liquid with pipette
    4. Repeat #2 3 more times
    5. Remove liquid

3. Permeabilization ( ii ) : 500-1000 µl washes
    1. Add PBT, Let sit for 10 minutes (3X)
    2. Remove PBT from well
4. Blocking : 500-1000µl washes
    1. Add 5% NGS in PBT (make personal aliquot and store in 4°C )
    2. Sit for 1 hr @ RT
5. Primary Incubation
    1. Dilute 1° in NGS in PBT in tubes, then administer 300-1000µl per well
        1. COMMONLY USED CTENO PRIMARIES dilutions
            1. Anti Acetylated Tubulin T6793 — use 1:400µL in 1.5mL tube
            2. Anti tyro tubulin - Millipore MAB1864 - use 1:1000
            3. mouse monocolonal hybridoma Bank typically 5ng/µl
    2. Cover dish and place inside of plastic plate — under darkness(foil wrapped) in the 4°C overnight
6. Wash ( i ) : 500-1000µl washes
    1. Wash out 1° antibody with 1mL of PBT for 10 mins (5X)
7. Secondary Incubation
    1. Dilute 2° Antibody in NGS in PBT
        1. 1:250
    2. Place 2° antibody solution in wells — 300-1000µl per well
    3. Cover dish with slide and inside of plastic plate — under darkness rocking in the 4°C
8. Wash ( iii ) : 500-1000µl washes
    1. Wash out 2° Antibody with 1mL of PBT for 10mins (5X) *** CRITICAL STEP***
9. Cell Staining
    1. DAPI (350 Blue)
        1. Use 1:1000
        2. Must mix solution in 1.5mL tube separate — centrifuge to homogenize — it will precipitate out if you put it directly on the sample
    2. Cell mask ( 488 Green)***optional***
        1. Use 1:1000
