## Supplementary figures and images for "*Notch* expression during ctenophore development gives insight into its ancestral function"

### SupFig1_Domain-architecture.jpg

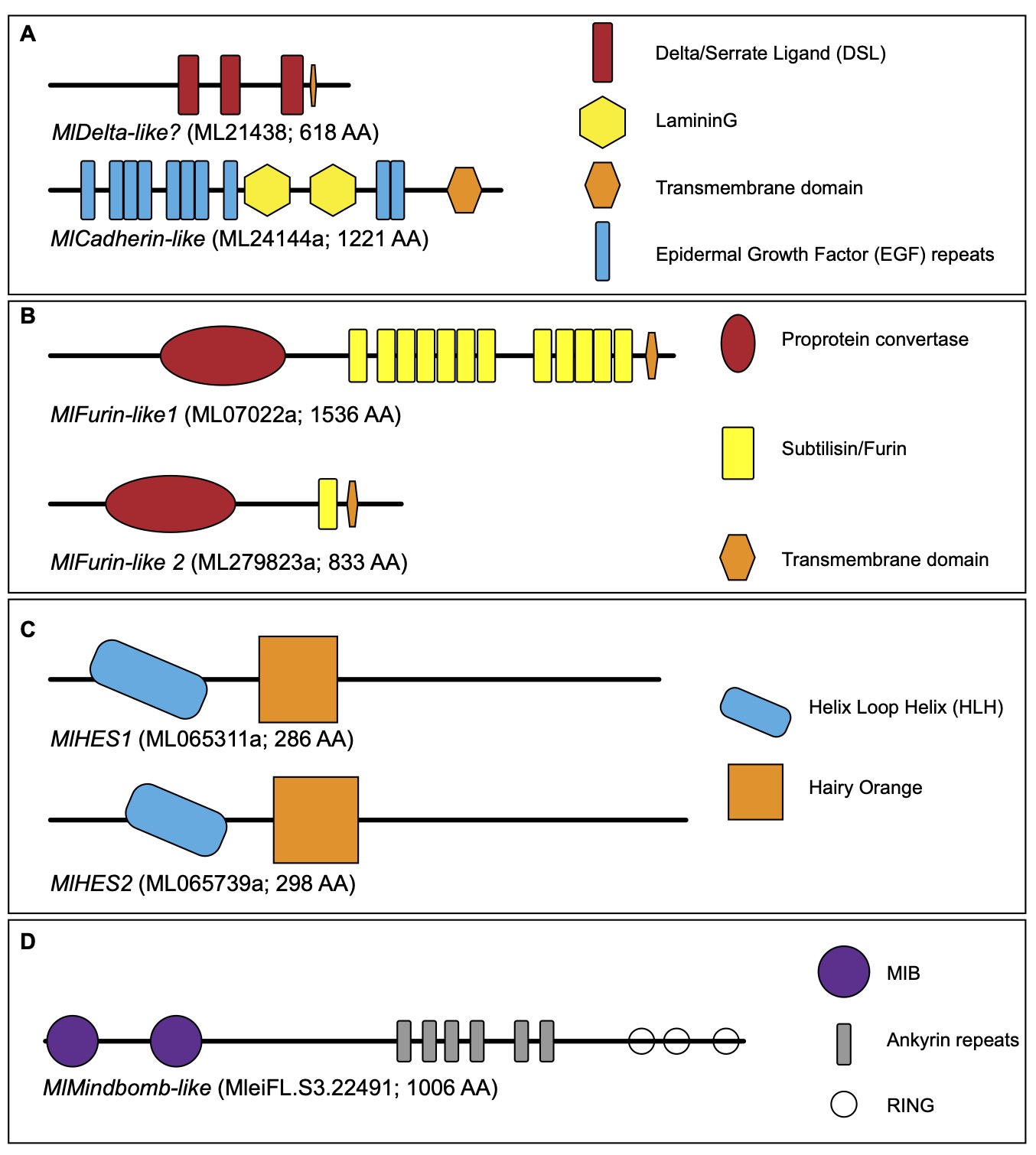

### SupFig2_MlDelta-like.jpg

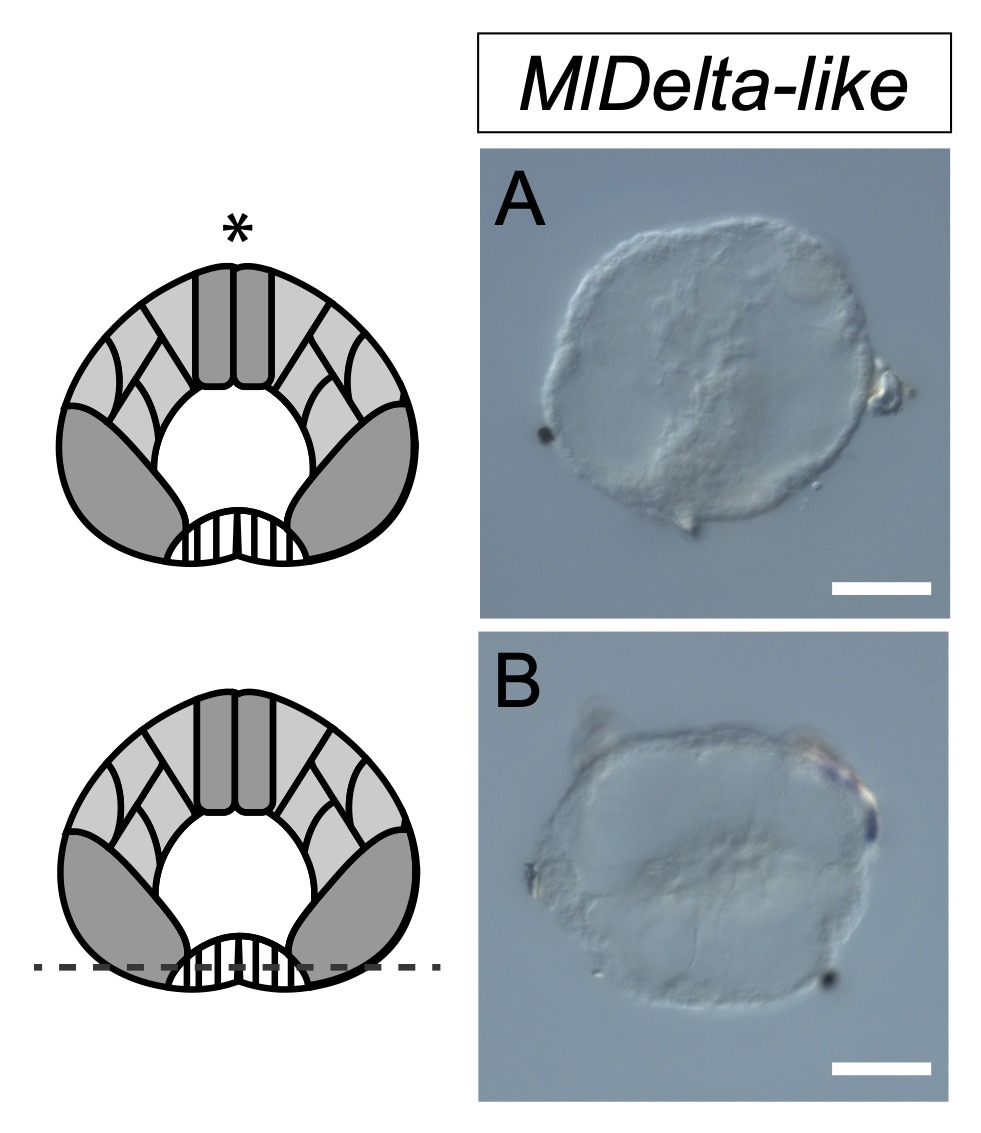

### SupFig3_Notch-24hpf.jpg

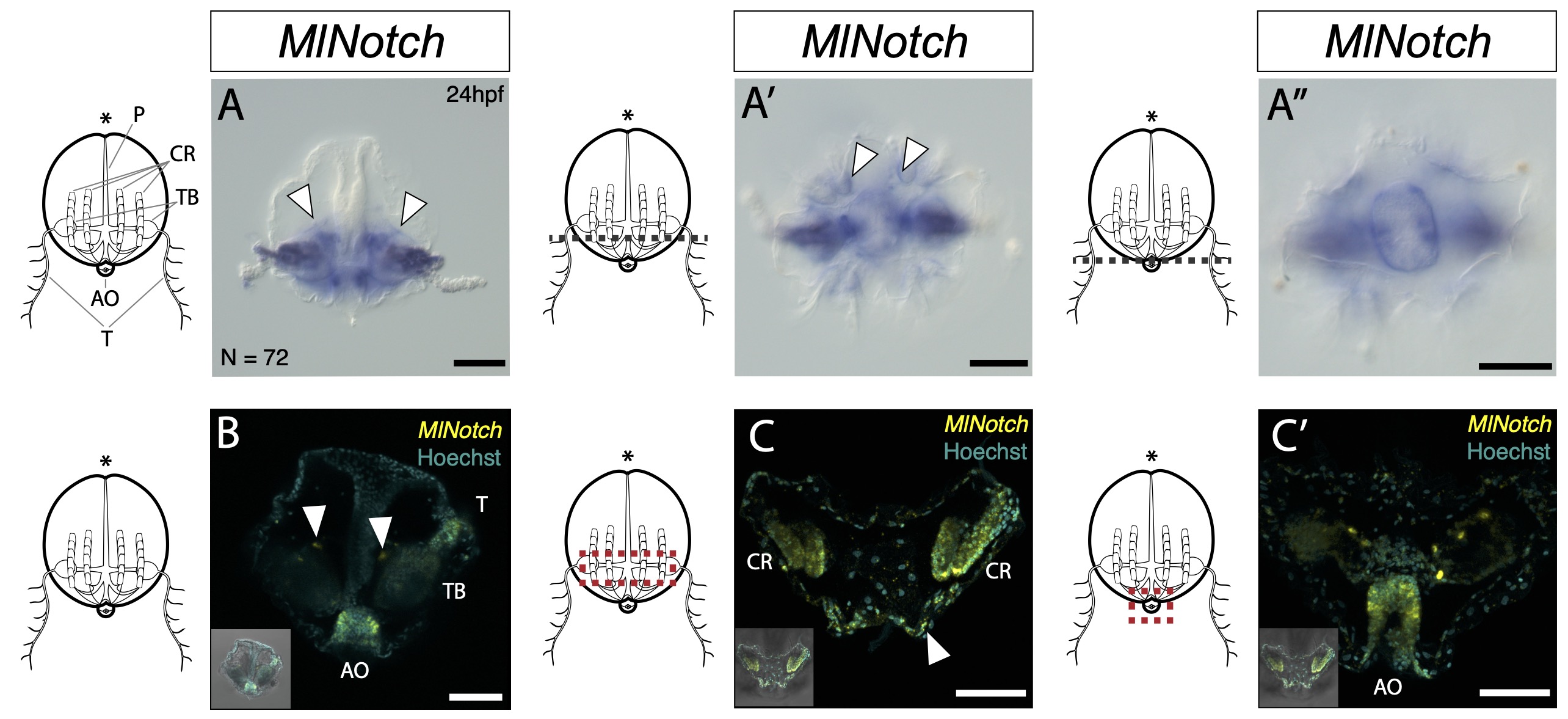

### SupFig4_cleaver-genes_24hpf.jpg

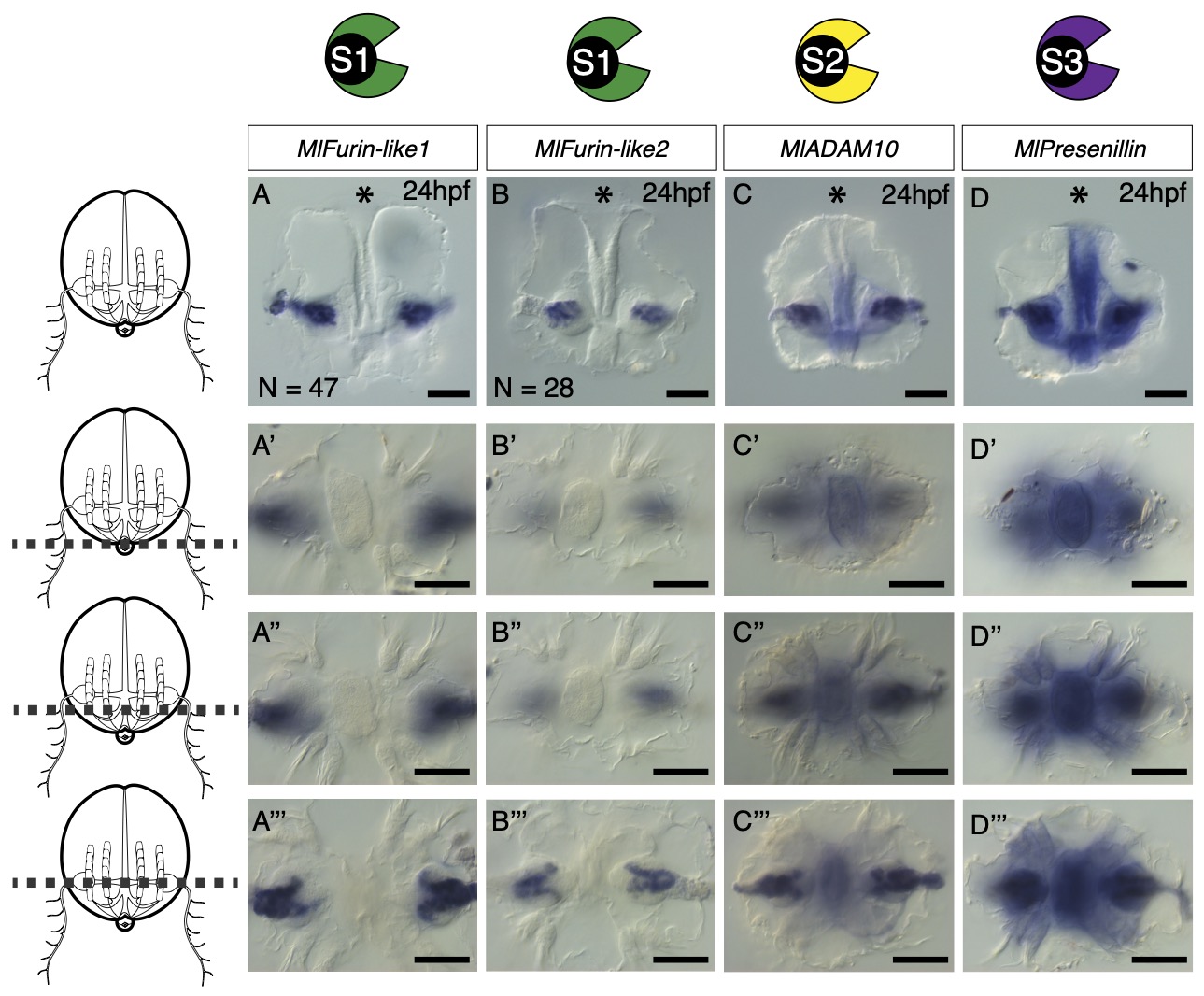
